# Effects of different environments and microbial exposures on the adult immune system of the C57BL/6 laboratory mouse strain

**DOI:** 10.1101/2025.10.22.683895

**Authors:** Kabeer Haneef, Meng Xie, Carlos F. Ibáñez

## Abstract

The realization that the grossly underdeveloped immune system of laboratory mice may hinder their use as suitable models of human disease has led to the development of different strategies to “normalize” immune responses in these mice so that they more closely resemble those typical of the human population. In this study, we performed a comprehensive comparison of the adult immune systems and microbiomes of C57BL/6 laboratory mice exposed to different housing and microbial conditions. C57/BL6 mice housed in a facility of substandard health levels (herein termed “dirty”) and C57BL/6 mice born after embryo transfer from females captured in the field (“wildings”) were compared to C57BL/6 mice housed under standard SPF laboratory conditions. Microbiota and parasite burden varied widely between the three conditions, which also showed marked differences among populations of mature, antigen-experienced, memory, plasma, and germinal centre B cells. Interestingly, we observed discrepancies among T cell phenotypes, with antigen-experienced T cells present at higher frequencies in “dirty” mice. On the other hand, “wildlings” displayed significantly elevated frequencies of both naïve and antigen-experienced spleen T cells. Interestingly, “dirty” and “wildlings” showed differences in the expansion of distinct myeloid cell types. Lastly, we observed that systemic immune traits reverted to a “cleaner” profile in the progenies (F2) of “wildlings” raised under laboratory conditions, indicating that such mice may not be suitable for human immune system modelling after prolonged housing in laboratory environments. Collectively, our results highlight the profound effects of varied microbial exposures and husbandry conditions on adult immunity of the common C57BL/6 laboratory mouse and suggest that complex microbial exposure in laboratory mice can provide a relevant tool for modelling immunological functions, thus enhancing their translational value.

## INTRODUCTION

Most of our understanding of the mammalian immune system is derived from genetically inbred and laboratory-adapted strains of the house mouse [1, 2], which are shielded from external events experienced by free-living mice, including complex social interactions, varied diets, weather fluctuations, and infections. Such an isolation from external variables significantly influences the immune system, contributing to the drastic differences found between the immune systems of adult laboratory mice and humans [3, 4]. Inbred laboratory mice housed in controlled conditions and carrying homogeneous gut microbiota allow a higher level of reproducibility in fundamental studies of mammalian development and physiology. As models of human disease, however, recent studies have indicated that their underdeveloped immune system may impose severe limitations on the utility of laboratory mice [5–8]. It has been suggested that free-living populations exposed to diverse environmental and microbial stimuli may more closely resemble the complexity of the adult human immune system and thus be better models for translational studies [2, 9, 10].On the other hand, the problem of such populations for biomedical research is their heterogenous genetic backgrounds.

Both the gut microbiome and the environment play crucial roles in numerous facets of human biology, regulating growth and development [11, 12], immune function [13, 14], metabolism [15], and subsequent immune protection against pathogenic insults [16]. Studies have shown that various environmental factors, including pollutants, nutrients, and social interaction, can alter the intestinal colonization of commensal microbiota, contributing to protection against hypersensitivity and autoimmune pathologies [17, 18]. In addition, the maternal microbiota has been reported to influence the development of foetuses and neonates through natural colonisation after birth, significantly reshaping the immune landscape of the offspring [19, 20]. A recent report showed that “wildlings,” derived by transferring laboratory mouse embryos into pseudopregnant feral female mice, inherit the complexity of the enteric gut microbiota from their foster mother and display a higher allergy-specific immune response upon exposure to HDM than laboratory mice [21]. Other studies have shown that cohousing with pet-store mice, which are typically kept under substandard sanitary conditions, significantly affects the immune system of inbred laboratory mice [9, 10], highlighting the significance of the effects of different environmental conditions on the immune landscape. Finally, some studies have exposed laboratory mice to a specific set of parasites [22, 23], or other natural conditions [10]. However, no study has so far made a direct comparison of these approaches and the question of whether different methods to “normalize” the immune system of laboratory mice achieve comparable outcomes remains unanswered.

In the present study, we have investigated the effects of two distinct approaches to “normalize” the immune system of C57BL/6 mice, a common strain of laboratory mice, on their cellular and humoral immune responses. To this end, two different models with varying degrees of microbial exposure and differences in husbandry conditions were established. Embryos from C57BL/6 mice housed in the laboratory under SPF (specific pathogen-free) conditions were transferred to mouse surrogate mothers captured in the wild, creating a distinct colony of “wilding” mice [24]. On the other hand, we obtained adult C57BL/6 mice from a facility that houses different kinds of rodents under substandard sanitary conditions, herein termed “dirty” mice. The microbiome, parasite burden and humoral and cellular components of the immune systems of “wildings” and “dirty” mice were studied in comparison to C57BL/6 mice housed under SPF conditions (herein referred to as “clean” or SPF mice). Although differences between the two models and laboratory “clean” mice were expected, we also found significant differences between the immune systems of “wildings” and “dirty” mice, suggesting that these two approaches, while offering clear advantages over clean SPF mice, may differentially model aspects of the human immune system. Our results will aid in designing suitable mouse models with broader clinical relevance for modelling human diseases and developing therapeutic strategies.

## RESULTS

### Generation of dirty and wildling mouse models to investigate the effects of gut microbiome complexity and parasite burden on immune system maturation

It is well accepted that humans are influenced by a variety of microbial encounters from early life, which play a key role in mediating immune development and function [25–27]. To better understand the effects of diverse microbial conditions that more closely resemble the state of the human immune system, we have generated two mouse models named “dirty” and **“**wildling”, respectively. “Dirty” C57BL/6 mice were procured from a rodent facility with substandard health levels (Fig. 1A). To generate wildlings, “clean” C57BL/6 embryos were transferred to feral female mice obtained from the field that were made pseudopregnant in the laboratory [24] (Fig. 1A). Cages housing weaned wildlings were supplemented with bedding from field mice to increase their microbial exposure during adolescence and kept on a different diet better resembling what they may encounter in the field (see Methods) [21]. Dirty mice and wildings were compared to mice raised in a clean environment under SPF conditions. Mice from the three groups were subjected to serology and parasitology analyses similar to those previously used in studies of mice from rewilding [10], pet stores [9] and wildlings [21]. The results are presented in Table S1. This analysis revealed divergent exposure to viral, bacterial, and helminth pathogens among the different mouse cohorts. In particular, dirty and field mice showed higher content of diverse pathogens and ectoparasites at epithelial barrier sites in comparison to wildlings, in agreement with the effects of diverse housing conditions on the occurrence and localization of different pathogens (Table S1).

**Figure 1.**
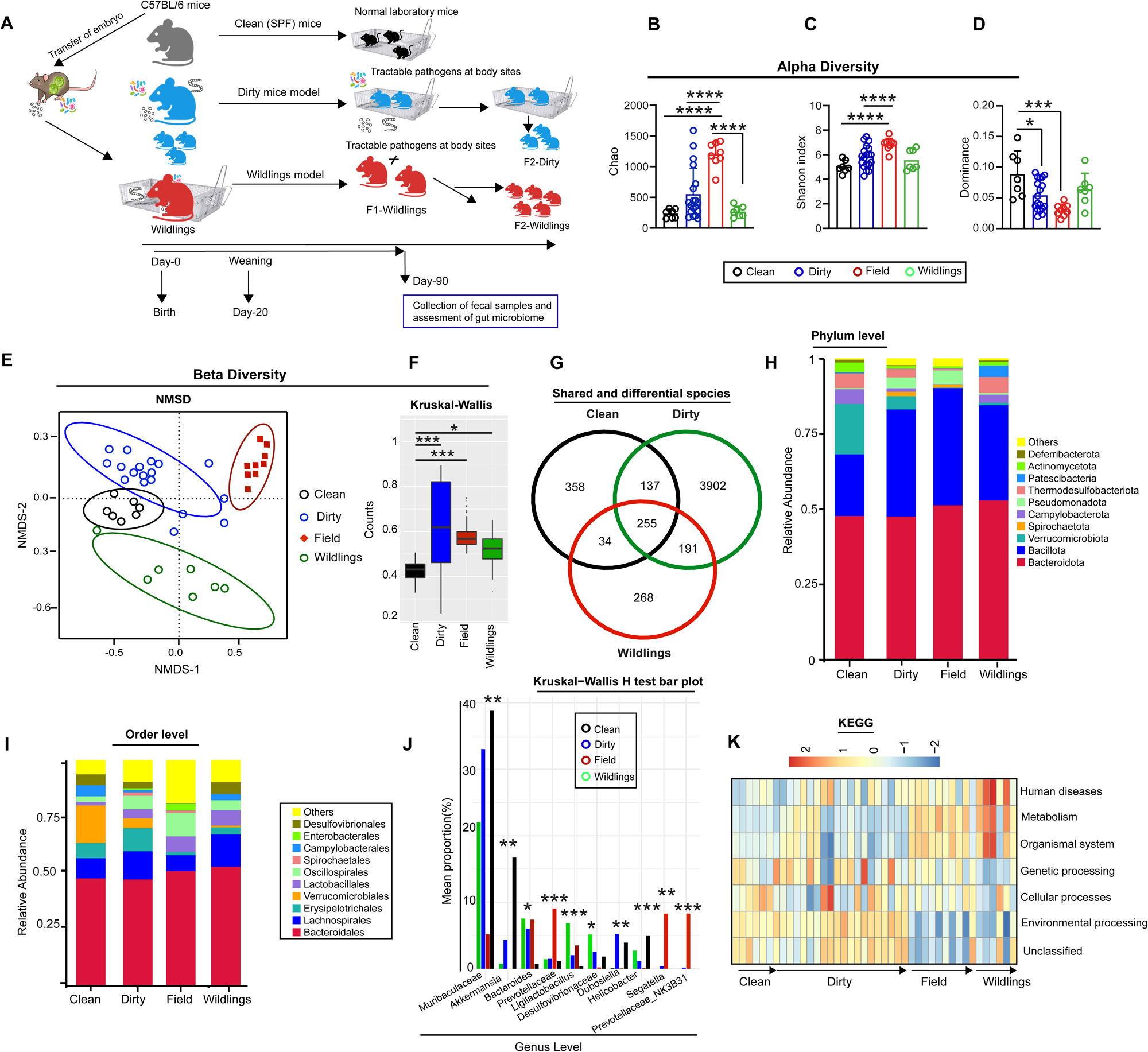
Generation of dirty and wildling mouse models to investigate the effects of gut microbiome complexity and parasite burden on immune system maturation. **(A)** The schematic figure shows different mouse models. ‘‘Dirty’’ mice were procured from a rodent facility with substandard health levels and ‘‘wildlings’’ were generated through transfer of “clean” embryos to pseudopregnant field mothers. Wildlings and dirty mice were self-crossed to acquire the next generations. **(B-D)** Bar plot displaying alpha diversity metrics, including Chao index **(B)**, Shannon index **(C)**, and Dominance analysis **(D)** among the gut microbiomes of all four groups**. (E)** NMDS analysis of the beta diversity of gut microbiomes among all four groups. Each dot represents a single mouse. Both axes indicate variations greater than 50%. **(F)** Kruskal-Wallis beta diversity box plot of variations in the microbiome composition among all groups. **(G)** Venn diagram displaying shared and distinct taxa among the gut microbiomes of clean, dirty, and wildlings. **(H, I)** Relative abundance of bacteria at phylum **(H)** and order **(I)** levels in the gut microbiome of clean, dirty, field and wildlings. **(J)** Differentially abundant taxa (Genus level) across clean, dirty, field, and wildling groups identified by Kruskal–Wallis H tests. Bars denote mean relative abundance (%) of each taxon per group. Bonferroni p-values are shown on the right (*, p < 0.05; **, p < 0.01; ***, p < 0.001). **(K)** Heatmap showing the relative enrichment (row-scaled Z-scores) of KEGG pathway categories across microbiomes from clean, dirty, field and wildling mice. Each column denotes an individual sample, while rows correspond to major KEGG functional categories across groups. Red indicates enrichment and blue indicates depletion relative to the mean. Dots in bar graphs correspond to individual mice. Means are shown. Statistical significance and *P* values were determined by non-parametric one-way ANOVA with Tukey’s multiple comparison tests.

It is well established that diverse environmental conditions can affect the composition of the gut microbiome [28, 29]. Thus, we first aimed to investigate the effects of the different environmental conditions on the diversity and enrichment of gut microbiota. To this end, 16S-RNA sequencing was conducted on fecal samples to survey the diversity of the enteric gut microbiome. Alpha diversity analysis, which measures the diversity within a single sample, revealed that field mice had a higher abundance of gut microbiome, as indicated by a higher Chao index (species richness) (Fig. 1B), compared to all other mouse cohorts. Similar patterns were observed in the Shannon index (richness and evenness) which was statistically significant in field mice compared to other groups (Fig. 1C). In contrast, dominance analysis revealed that clean mice have a relatively higher abundance of species that dominate the composition of the gut microbiome compared to other groups (Fig. 1D). Next, we applied non-metric multidimensional scaling (NMSD) using Bray Curtis dissimilarity analysis to examine beta diversity, which compares the similarity among different samples, and noticed significant variations among the gut microbiota of all groups (Fig. 1E). Interestingly, the cluster corresponding to wildling mice microbiome exhibited substantial divergence from that of field mice, showing a trend towards convergence with the microbiome of clean SPF-raised mice (Fig. 1E), an effect that may be due to prolonged husbandry in a laboratory (albeit non-SPF) environment. Further beta-diversity analysis, as shown in the Kruskal-Wallis plot and Venn diagram, revealed significant variations among the gut microbiomes of all groups (Fig. 1F, G).

Taxonomic evaluation revealed that, at the phylum level, dirty, wildlings and field cohorts displayed increased abundance of *Bacteroidota* and *Bacillota* (phylum level), compared to clean mice (Fig. 1H), the latter increase at the expense of *Verrucomicrobiota*, which was only observed at significant proportions in clean mice (Fig. 1H). At the order level, clean mice displayed an increased abundance of *Verrucomicrobiales* compared to the other groups (Fig. 1I). Interestingly, *Bacteriodales* (order level) was detected at significantly higher levels in field mice and wildlings (Fig. 1I), a feature that was likely transferred from field mothers to their wildling offspring. Next, we examined taxonomic differences in the gut microbiome across all cohorts using Kruskal–Wallis H tests (FDR-corrected), which revealed significant variations among the four groups (Fig. 1J). *Muribaculaceae* and *Akkermansia* (Genus level) were significantly enriched in clean mice, whereas *Prevotellaceae*, *Helicobacter, Ligiliatobacillus*, and *Prevotellaceae_NK3B31*_group were substantially more abundant in the field and wildling groups. In contrast, dirty mice were characterized by higher levels of *Segatella* and *Dubosiella*, supporting the effects of different environments on the diversity of the gut microbiome (Fig. 1J). We further assessed the functional enrichment of gut microbiome across all mouse cohorts using KEGG pathway annotations, revealing subtle differences (Fig. 1K). Notably, the clean group was enriched in cellular pathways that support fundamental cell-level activities rather than metabolism. In contrast, the dirty group showed significant microbiome enrichment in cellular functions and genetic information processing pathways that support motility, cell development, death and protein translation [32] (Fig. 1K). In contrast, wildlings possessed a microbiome that was functionally unique from the other groups and possessed pathways associated with human diseases (including infectious and immunological-related processes), metabolism (carbohydrate and lipid metabolism), and organismal systems (digestive, immune and endocrine modules), supporting the notion that wildling microbiota is both metabolically adaptable and functionally diverse towards microbial and pathogenic insults in their natural environment. In line with this, the field mouse microbiome exhibited enrichment in pathways associated with metabolism and organismal systems (Fig. 1K).

Lastly, the possible effects of prolonged husbandry in a laboratory environment on the composition of the gut microbiome were investigated. Specifically, we compared the microbiome of newly procured field mice to that of field mice kept at the non-SPF section of the laboratory vivarium for 8 weeks. Principal coordinate analysis (PCoA), alpha diversity and taxon analysis revealed notable variations among the gut microbiota, suggesting that prolonged captivity of field mice in a laboratory environment significantly alters the composition of the gut microbiome (Fig. S1, A-I). Overall, these findings indicate that diverse microbial exposures and environmental conditions strongly affect the composition of the gut microbiome. Subsequent analyses of immune cell composition and function focused on C57BL/6 cohorts of clean, dirty, and wildling mice, given that field mice are outbred and of unknown genetic composition.

### Distinct effects of altered microbial exposures and housing conditions on the expansion of B cells

There is an increasing appreciation that microbial exposure and gut microbiota can significantly impact the development and functioning of the immune system [13, 14, 33, 34]. To investigate the impact of different microbial exposures and housing conditions on B cell development and maturation in peripheral and regional lymphoid organs, we first examined spleens from clean, dirty, and wildling mouse cohorts (Fig. 2A). Evaluation of the spleen index at postnatal day 90 (P90) revealed that dirty mice had significantly increased spleen weight (*P=0.001*) compared to clean and wildling cohorts (Fig. 2B). Next, to examine B cell development and expansion, we quantified B220 (CD45R) and CD19 expressing cells to assess mature and resting B cells, respectively. The loss of CD19 expression is characteristic of differentiated plasma B cells [35]. Therefore, CD19^−^B220^+^ population in peripheral circulation and regional lymphoid organs may represent differentiated B cells [36, 37], while the presence of CD19 characterizes developing B cells [38] (Fig. S2A, B). Flow cytometry revealed that both dirty and wildlings exhibited significantly higher percentages (p <0.001) of differentiated (CD19-B220+) spleen B cells compared to SPF-clean mice (Fig. 2C). In addition, we observed that only wildlings had significantly higher proportions of mature (CD19+B220+) B cells (Fig. 2D). In contrast, dirty mice exhibited significantly lower frequencies of mature B cells than clean and wildlings (Fig. 2D), supporting the notion that diverse microbial exposure and altered microbiota can affect B cell maturation.

**Figure 2.**
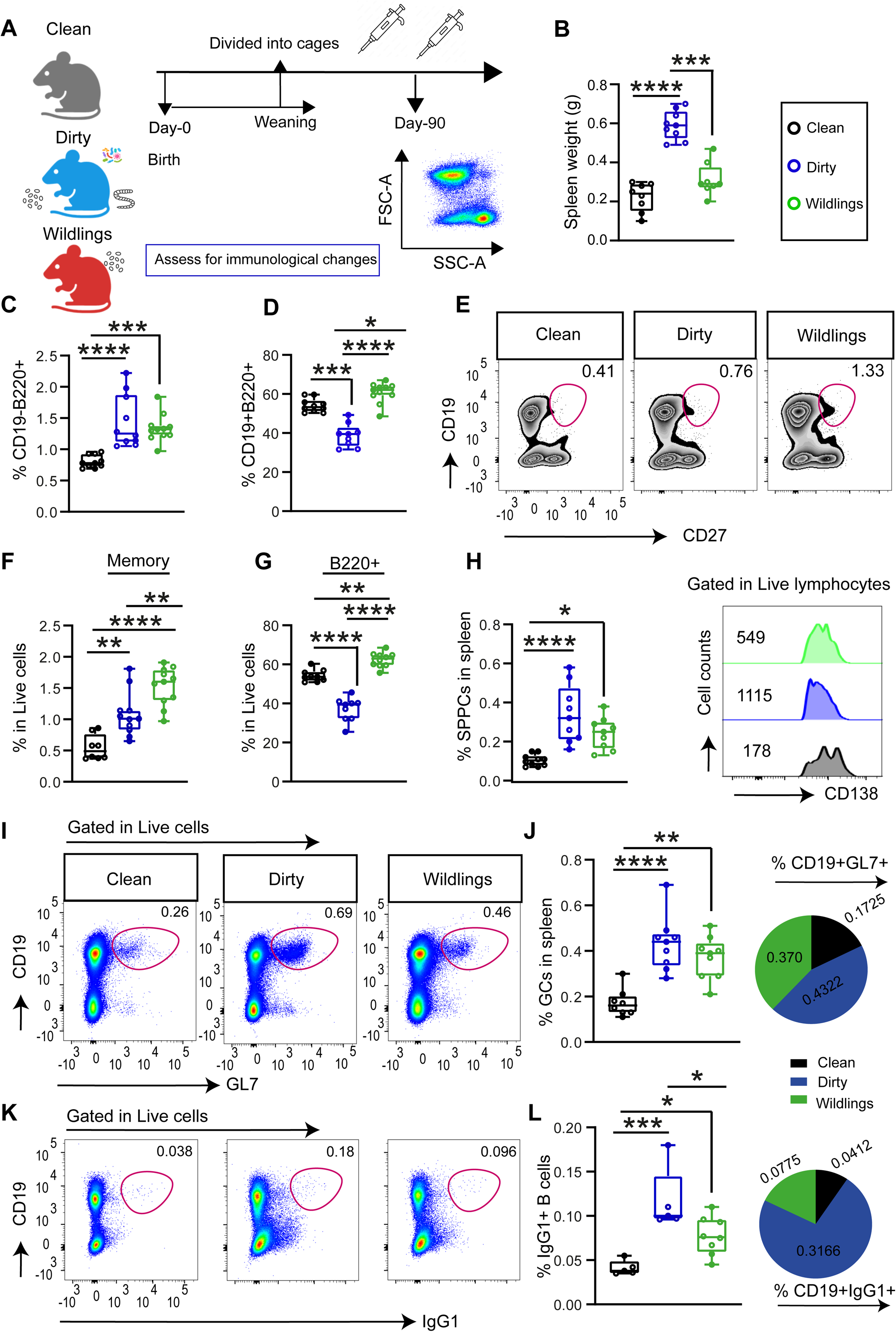
Distinct effects of altered microbial exposures and housing conditions on the expansion of B cells. **(A)** Schematic showing clean (SPF), dirty, and wildlings mice models. Mice were divided into cages after weaning, and the spleen of clean, dirty and wildlings was collected at postnatal-90 (p-90) for flow cytometry analysis. **(B)** measurement of spleen index from clean, dirty and wildlings. **(C, D)** Frequencies of differentiated **(C)** and mature **(D)** B cells in the spleen of clean, dirty and wildlings. **(E)** Representative flow cytometry contour plots of CD19 versus CD27 expression in the spleen of clean, dirty and wildlings. **(F)** Frequencies of memory B cell subsets (CD19+CD27+) from **E**. **(G, H)** Summary statistic plots show the frequency of B220+ B cells **(G)** and antibody-secreting plasma cells (B220-CD138+) in the spleen of all 3 mouse cohorts. **(I)** Representative flow cytometry pseudocolour plots of CD19 versus GL7 expression in the spleen of clean, dirty and wildlings. **(J)** Frequencies of germinal center B cells (CD19+GL7+) from **I**, and the pie chart (right) shows the mean distribution frequencies of GCS in the spleen of clean, dirty, and wildlings. (**K, L)** Representative flow cytometry pseudocolour plots of CD19 versus IgG1 expression **(K)** and frequencies of IgG1+ B cells (middle) from **(K),** and pie chart (right) shows the average distribution percentages of isotype class-switched IgG1+ B cells in the spleen of clean, dirty and wildling cohorts. Dots in bar graphs correspond to individual mice. Means are shown. Statistical significance and *P* values were determined by non-parametric one-way ANOVA with Tukey’s multiple comparison tests; *p < 0.05; **p < 0.01.

Next, we assessed the pool of plasma and memory B cells, which are terminally differentiated immune cell subsets that provide a defensive response against pathogens [35, 39]. Since stimulating environments induce the differentiation of B cells into antibody-secreting plasma and memory B cell subsets [40, 41], we hypothesized that mice with diverse microbial exposure would display higher frequencies of plasma and memory B cell subsets. Flow cytometric profiling showed that dirty and wildling mice had increased frequencies of memory (CD19+CD27^+^) B cells (Fig. 2E, F). Notably, the spleen of wildlings exhibited an expanded population of memory B cells compared to dirty and clean mice (Fig. 2E, F). Interestingly, dirty mice showed a significantly lower frequency of B220+ B cells than the clean and wildling cohorts (Fig. 2G). As expected, we observed increased frequencies of spleen plasma cells (SPPCs) (CD138+B220-) in both dirty and wildling mice compared to clean mice (Fig. 2H).

Germinal center (GC) B cells play a crucial role in affinity maturation and generation of high-affinity antibodies [41, 42]. To assess GC responses, we collected spleens and examined the expression of GL7 using flow cytometry. GL7 is upregulated in activated B cells, but not in naïve B cells, and is widely used as a marker for germinal center B cells in secondary lymphoid organs [42]. As expected, we observed that the spleens of dirty and wildlings contained significantly higher frequencies of GC B cells (CD19+GL7+) relative to clean mice (Fig. 2I, J). Consistent with the significant increase in GC B cells, dirty and wildlings exhibited higher percentages of class-switched IgG1+ and IgG2b+ B cells in the spleen (Fig. 2K, L, and Fig. S2C). Of note, the degree of GC responses and isotype class-switching was more modest in dirty mice than what was observed in the spleen of wildlings. These findings demonstrate enhanced antigen-experienced humoral immune responses in dirty mice compared to the other groups. Overall, our analysis of B cell populations demonstrate that diversity of environmental conditions induce differential B cell profiles.

### Effects of altered microbial exposures and housing conditions on spleen T cell pools

Next, we sought to understand how diverse microbial exposures and environmental perturbations affect naïve and antigen-experienced T cells in the spleen. T cells are crucial for cell-mediated immune responses towards intracellular pathogens, and their proximal expansion and distribution directly relate to their function [43–45]. Single-cell spleen suspensions were analyzed using flow cytometry and t-distributed stochastic neighbor embedding (t-SNE) visualization to assess the distribution of immune cells. We evaluated the spleen, as the gut-spleen axis facilitates communication between multiple organ systems [46]. Flow cytometry analysis revealed disparities in the frequencies of naïve (CD8+CD4-, CD4+CD8-, CD4+CD62L+) and antigen-experienced virtual memory (TVM, CD44+CD49d-of CD8+CD4-) and true memory (TTM, CD44+CD49d+ of CD8+CD4-) T cell subtypes. Splenic CD8+CD4- and CD8-CD4+ T cells were present at increased frequencies in the spleens of wildlings; while dirty mice exhibited a significantly lower proportion of these cells in comparison to clean and wildling cohorts (Fig. 3A, B), indicating the effects of differential environmental exposures on the composition of T cells in the spleen. To gain further insights, we conducted a comprehensive phenotypic assessment of differentiation markers previously identified in CD8+ T cells (Fig. S3A). This revealed a significant expansion of antigen-experienced ‘CD8-TVM’ and ‘CD8-TTM-T’ cells in the spleens of dirty mice (Fig. 3C, D, E). Notably, the data showed that only the spleens of dirty mice exhibited increased frequencies of both types of antigen-experienced T cells (CD8-TTM and TVM) compared to wildlings and clean mice (Fig. 3C, D, E). This could be due to the particular set of pathogens, ectoparasites, and microbial composition that characterizes this group of mice. In contrast, the spleen of wildling mice showed frequencies of CD8-TVM that were comparable to clean mice (Fig. 3C, D), but displayed elevated percentages of true memory (CD8+TTM) T cells (Fig. 3C, E). Moreover, the frequency of CD8-memory T cells (CD8+CD44+) was increased in the spleens of dirty mice and wildlings (Fig. 3F, G, and Fig. S3B). In line with elevated levels of antigen-experienced T cells, we also quantified the expression of costimulatory molecules, which are essential for T-B cell interaction, including CD62L [47], and observed a marked reduction in the frequencies of CD4+62L+ T cells in the spleens of dirty mice compared to clean and wildlings (Fig. 3H, I). We further assessed the distribution of different T cell subsets using t-SNE visualization which confirmed the disparity among the T cell phenotypes of the three groups of mice (Fig. 3J). These results indicate that diverse environmental exposures induce differential residency of activated T cells, enhancing their activated phenotypes, which in turn may facilitate B cell activation and differentiation.

**Figure 3:**
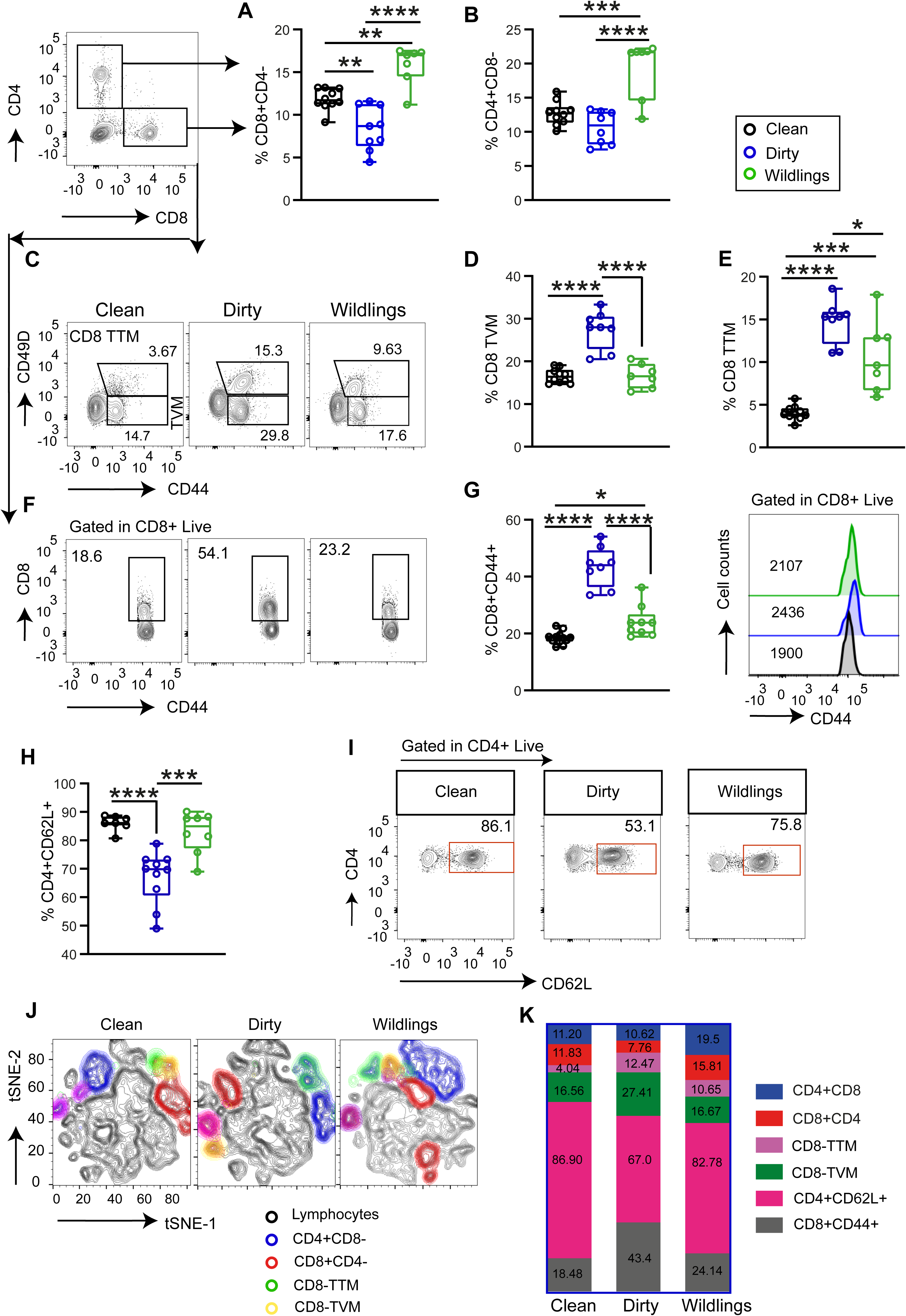
Effects of altered microbial exposures and housing conditions on spleen T cell pools. **(A, B)** Frequencies of CD8+CD4- **(A),** CD4+CD8- **(B)** T cells from the spleens of clean, dirty, and wildling mice. **(C)** Representative flow cytometry contour plots showing the expression of CD44 versus CD49d in gated CD8+CD4-T cells from the spleens of clean, dirty and wildling mice. **(D, E)** Frequencies of CD8 TVM (CD44+CD49d-of CD8+CD4-) and CD8 TTM (CD44+CD49d+ of CD8+CD4-) T cells gated from the CD8+CD4-T cells in the spleen of clean, dirty and wildling cohorts. **(F, G)** Representative flow cytometry contour plots showing the expression of CD8 versus CD44 gated from CD8+ T cells **(F),** and summary statistic plot shows the corresponding frequencies of CD8+CD44+ **(G)**, histogram plot (right) shows the expression of CD44+ cells in gated CD8+CD44+ T cells in the spleens of clean, dirty, and wildlings mice. **(H, I)** summary statistic plot shows the frequencies of CD4+CD62L+ T cells **(H),** and representative flow cytometry contour plots (right) showing the expression of CD4 versus CD62L gated from CD4+ T cells**(H),** in the spleen of clean, dirty and wildlings. **(J)** t-SNE plot displays the distribution of different naïve and antigen-experienced T cell subsets from the spleen of these three mouse cohorts. **(K)** Mean distribution frequencies of different naïve and antigen-experienced T cell subsets in the spleen of clean, dirty and wildlings. Each colour represents the cell type. Each dot represents a single mouse. Statistical significance and *P* values were determined by non-parametric one-way ANOVA with Tukey’s multiple comparison tests; *p < 0.05; **p < 0.01.

### Effects of environmental conditions on myeloid cells

Next, we measured the effect of altered microbial and housing conditions on the expansion of myeloid cells in the spleen using a combination of markers (Fig. 4A). Myeloid cells are key components of the innate immune system, serving as an essential interface between the innate and adaptive immune systems [48]. We assessed the expansion of different myeloid cell populations in the spleen by flow cytometry as illustrated in Fig. S4A. High side-scatter (SSC^high^) high forward scatter (FSC^high^) cells typically characterize populations of granulocytes. As expected, we observed that spleens from dirty mice displayed increased frequencies of these cells (SSC^high^FSC^high^) compared to the clean and wildling cohorts (Fig. 4B). We further observed that dirty mice had significantly higher percentages of CD11b+ cells (Fig. 4C), a marker of myeloid lineages including macrophages, neutrophils, and subsets of dendritic cells [49, 50], while clean and wildings did not differ with regards to these cell populations (Fig. 4C).

**Figure 4.**
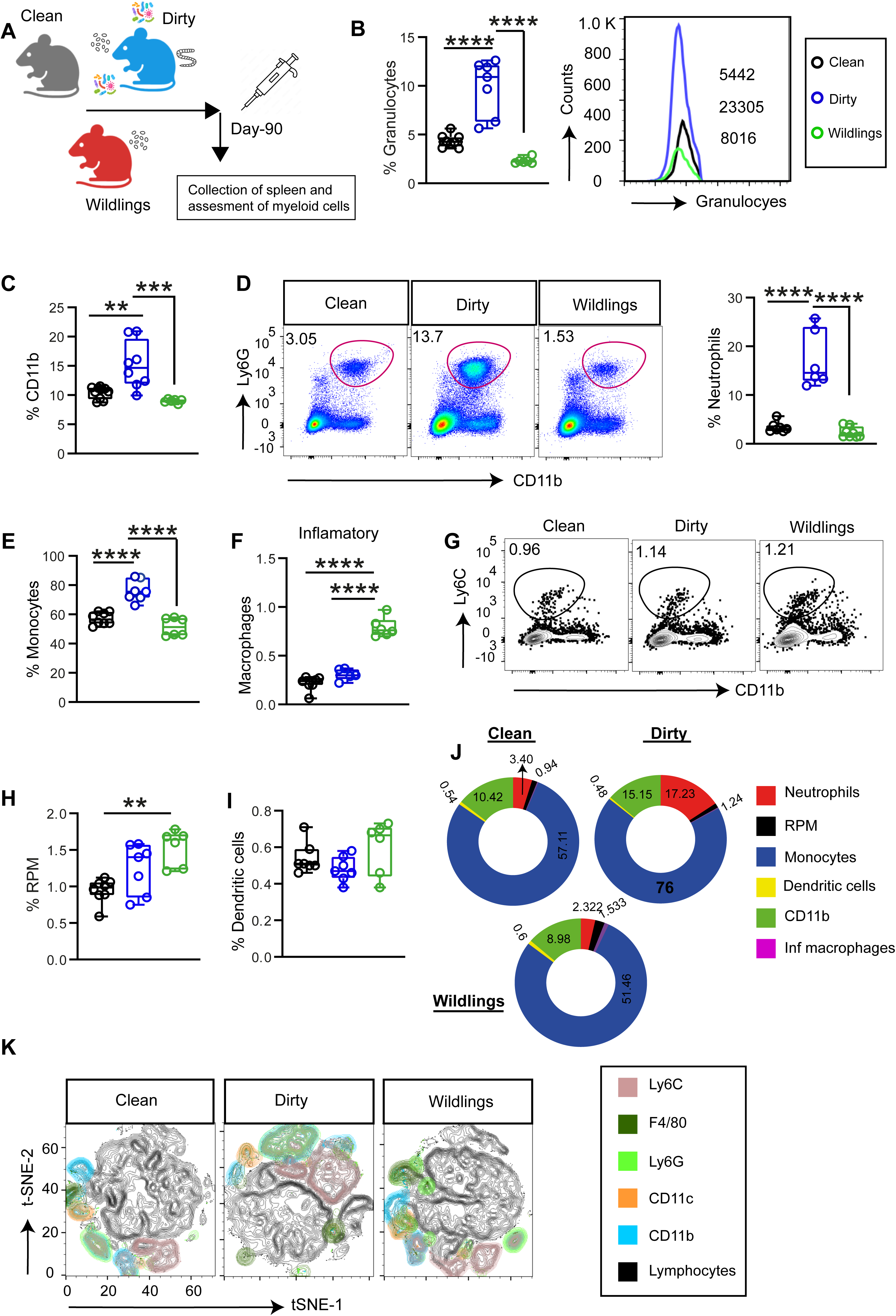
Effects of environmental conditions on myeloid cells. **(A)** Schematic of clean, dirty, and wildling mice. 3-week-old mice were sacrificed at p-90 for flow cytometry analysis. **(B)** Bar plot depicts the frequencies of granulocytes (SSC^high^FSC^high^) (middle), and the histogram on the right shows the relative granulocyte cell counts. **(C)** Quantification of the frequencies of CD11b+ cells. **(D)** Representative flow cytometry pseudocolour plots showing the expression of Ly6G versus CD11b gated from DAPI-live cells (middle) and bar graph (right) displaying the frequencies of neutrophils (CD11b+Ly6G+) in the spleens of clean, dirty and wildlings. **(E, F)** Frequencies of monocytes **(E)** and macrophages **(F)** in the spleens of clean, dirty and wildlings. **(G)** Representative flow cytometry contour plots display the expression of Ly6c versus CD11b in the spleens of clean, dirty and wildlings. **(H, I)** Summary statistics of the frequencies of red pulp macrophages (F4/80^high^CD11b^low^) **(H)** and dendritic cells (CD11b^high^CD11c^high^) **(I)** in the spleen of all three mouse cohorts. **(J)** Pie charts illustrate the mean distribution frequencies of different myeloid cells in clean, dirty, and wildling spleens. **(K)** Different myeloid subsets were quantified by flow cytometry from the spleens and visualized with two-dimensional t-SNE. Each dot represents a single mouse. Data are represented as mean SEM. Statistical significance and *P* values were determined by non-parametric one-way ANOVA with Tukey’s multiple comparison tests; *p < 0.05; **p < 0.01.

The lymphocyte antigen-6 complex (Ly6G) and CD11b were used to distinguish neutrophils from other myeloid cells [51]. Again, the spleens of dirty mice displayed elevated frequencies of neutrophils (CD11b+Ly6G+) compared to the other two groups (Fig. 4D). In line with the marked expansion of neutrophils, dirty mice spleens displayed increased frequencies of monocytes (Ly6c^high^Ly6c^low^) compared to both clean and wildling counterparts (Fig. 4E). In contrast, inflammatory macrophages (CD11b+Ly6c+F4/80+) (Fig. 4F) and tissue resident red-pulp macrophages (F4/80^high^CD11b^low^) (Fig. 4G, H) were present at higher frequencies in the spleen of wildlings, while clean and dirty mice were comparable in this regard (Fig. 4F, G, H), indicating differential expansion of diverse classes of myeloid cells in response to altered microbial exposures and housing conditions. However, spleen dendritic cells (CD11b^high^CD11c^high^) were comparable across all three mouse cohorts (Fig. 4, I). Further analysis of myeloid markers through t-SNE demonstrated notable differences among the expansion of different myeloid-specific markers (CD11b, Ly6G, F4/80, CD11c) in spleens of dirty mice compared to clean mice and wildlings (Fig. 4K).

To further investigate whether these observations are feature exclusive to the spleen or extend to other regions, we analyzed the frequencies of different classes of myeloid cells in the peripheral circulation. Compared to both clean mice and wildlings, dirty mice exhibited increased percentages of circulating granulocytes (Fig. S4C), in agreement with our observations in spleen. On the other hand, higher frequencies of circulating dendritic cells were found in the blood of wildlings compared to the other two groups (Fig. S4D). We also quantified the frequencies of monocytes and CD11b+ myeloid cells in the mesenteric lymph nodes (MLN) and observed higher percentages of CD11b+ myeloid cells (Fig. S4E), and monocytes (Fig. S4F, G) in the MLN of dirty mice as compared to clean and wildling cohorts. Overall, these findings demonstrate the impact of microbial diversity and altered husbandry conditions on the composition and diversity of myeloid cells in peripheral and regional lymphoid organs.

### The immune phenotype of wildlings converges to that of clean mice after subsequent breeding in a laboratory environment

It has been reported that wildlings exhibit stability of microbiota and preserve the characteristic activation of the immune system of their wild counterparts [24]. However, other results have shown that long-term environmental exposure and altered dietary patterns can significantly affect the gut microbiota and influence immune responses [34, 52]. Therefore, we addressed the question of whether second-generation (F2) wildlings born and raised in a laboratory environment (albeit non-SPF) possessed the functional systemic immune traits of F1 wildlings born to field mice (Fig. 5A). Pathogen assessment revealed a significant reduction in the proportions of ectoparasites in F2 wildlings compared to F1 wildlings (Table 1), suggesting that laboratory breeding can reduce pathogen colonization. We then compared the frequencies of several circulating immune cell subsets. Both F1 and F2 wildlings exhibited comparable proportions of circulating B220+CD3-cells (Fig. 5B), while F2 displayed a marked reduction in the frequencies of CD3+B220-T cells (Fig. 5C). In addition, we further observed that F2 wildlings showed a significant decrease in the frequencies of memory (CD19+CD27+) (Fig. 5D) and circulating plasma B cells subsets (CD138+) compared to F1 wildlings (Fig. 5 E). Intriguingly, however, neutrophils were present at comparable frequencies in both F1 and F2 wilding cohorts (Fig. 5F). With regards to circulating T cells, F2 wildlings displayed a notable reduction in the frequencies of both antigen-experienced CD8+ TTM and CD8+ TVM T cells (Fig. 5H) compared to the F1 generation. Interestingly, naïve T cells (CD8+, CD4+) were present at significantly higher percentages in the blood of F2 wildlings as compared to F1 (Fig. S5A, B). In contrast, F1 and F2 wildlings were comparable in the expression of CD44+ cells, a marker associated with T-cell activation, migration and immune memory [53, 54] (Fig. 5I). Further phenotypic assessment revealed a significant decrease in the frequencies of CD4+CD44+ (Fig. 5J) and CD8+CD44+ T cells in F2 wildlings (Fig. 5K), indicating that prolonged housing of wildlings in a laboratory environment alters the maturation of systemic immune system properties. For further validation, we quantified the frequencies of distinct sets of myeloid cells in the F2 progenies of ‘‘dirty mice’’ generated in the laboratory environment (Fig. S5C), and observed that F2 dirty mice showed significantly lower percentages of circulating granulocytes (Fig. S5D) and monocytes (Fig. S5E) compared to the F1 generation. We further observed the significant decrease in the percentages of neutrophils (Fig. S5F) and CD11b+ myeloid cells (Fig. S5G) in F2 progenies of ‘‘dirty mice’’. Intriguingly, dendritic cells were present at comparable frequencies in both F1 and F2 generations of dirty mice (Fig. S5G). Overall, these findings indicate that F1 wildlings exhibit enhanced activation of the immune system compared to F2, suggesting that care should be exercised when breeding populations of wildlings (or other types of “dirty” mice) in the laboratory for downstream studies of immune cell composition and activity.

**Figure 5.**
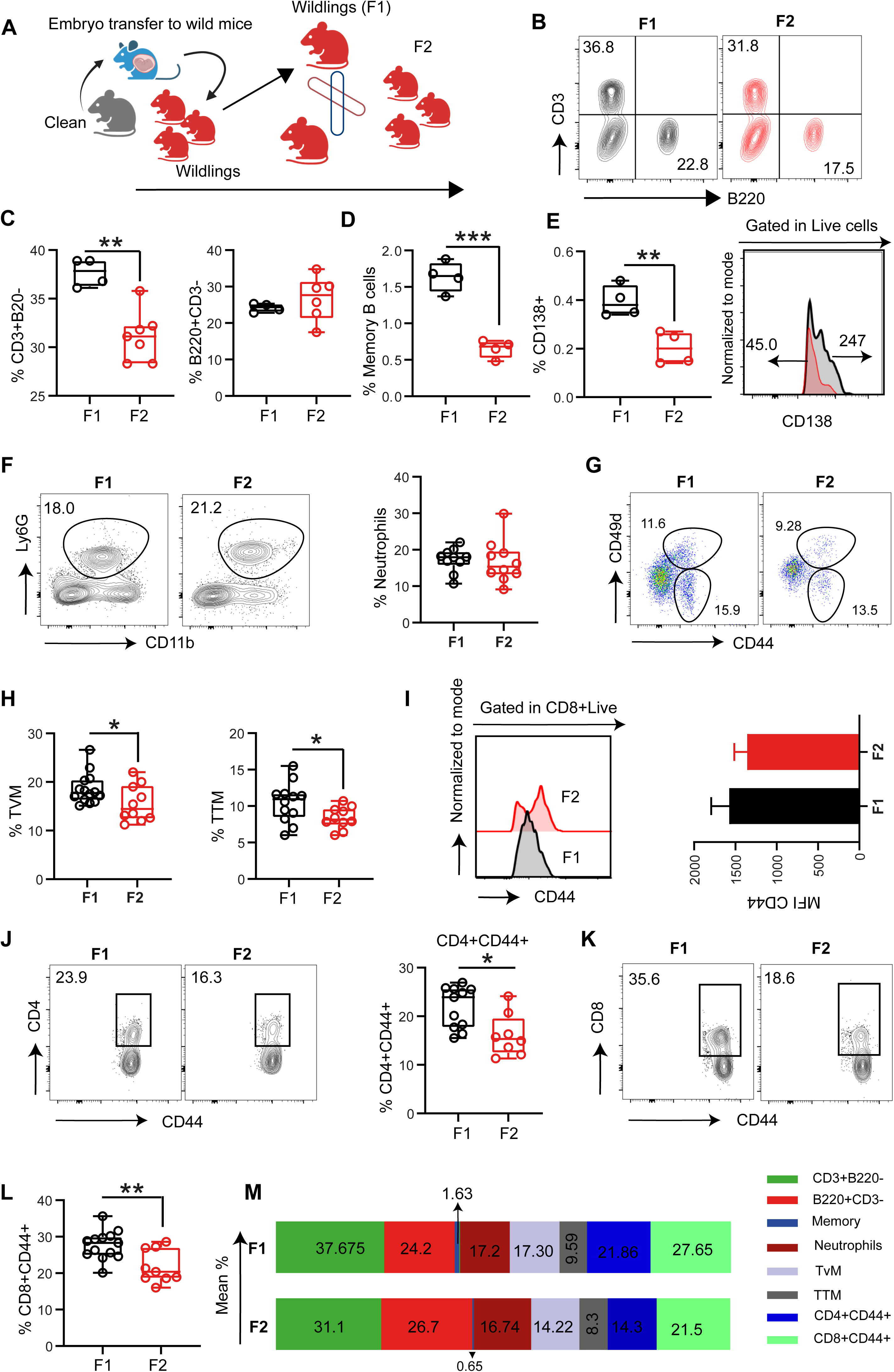
The immune phenotypes of wildlings converge to those of clean mice after two generations in a laboratory environment. **(A)** Schematic of F1 and F2 wildlings and comparative assessment of immune phenotypes at postnatal day 90 from the peripheral circulation. **(B)** Representative flow cytometry contour plots showing the B220 versus CD3 expression (top right) in the blood of F1 and F2 wildlings. **(C)** Summary statistic plot showing the quantification of the frequencies of CD3+B220- and B220+CD3-from B. **(D)** Frequencies of memory B cell (CD19+CD2+) in the peripheral circulation of F1 and F2 wildlings. **(E)** Summary statistic plot displays the frequencies of CD138+ cells, and histogram (right) shows the relative cell counts from the blood of F1 and F2 wildlings. **(F)** Representative flow cytometry contour plots displaying the Ly6G versus CD11b expression from the peripheral circulation of F1 and F2 wildlings (left), and summary statistic graph (middle) shows the quantification of the frequencies of neutrophils (Ly6G+CD11b+) from F. **(G)** Flow cytometry pseudocolour plots showing the CD44 versus CD49d expression gated in CD8+CD4-live cells from the peripheral circulation of F1 and F2 wildlings. **(H)** Frequencies of different antigen-experienced T cells, specifically CD8-TVM and CD8-TTM, in G. **(I)** A representative histogram plot illustrates the expression of CD44+ cells in gated CD44+CD49D+ cells, and a corresponding bar graph (right) displays the mean fluorescence intensity (MFI) values in the blood of F1 and F2 wildlings. **(J)** Representative flow cytometry contour plots displaying the CD4 versus CD44 expression (left) and a summary statistics plot (middle) show the quantification of the frequencies of CD4+CD44+ T cells from J. **(K)** Representative flow cytometry contour plots show the CD8 versus CD44 expression (right) in the blood of F1 and F2 wildlings. **(L)** Summary statistics of the frequencies of CD8+CD44+ T cells from K. **(M)** Bar graphs displaying the mean distribution frequencies of different circulating immune cells in the blood of F1 and F2 wildlings. Each colour represents the cell type. Data are represented as mean SD. Statistical evaluation was done using an unpaired two-tailed Student t-test; *p < 0.05; **p < 0.01.

**Table 1.**
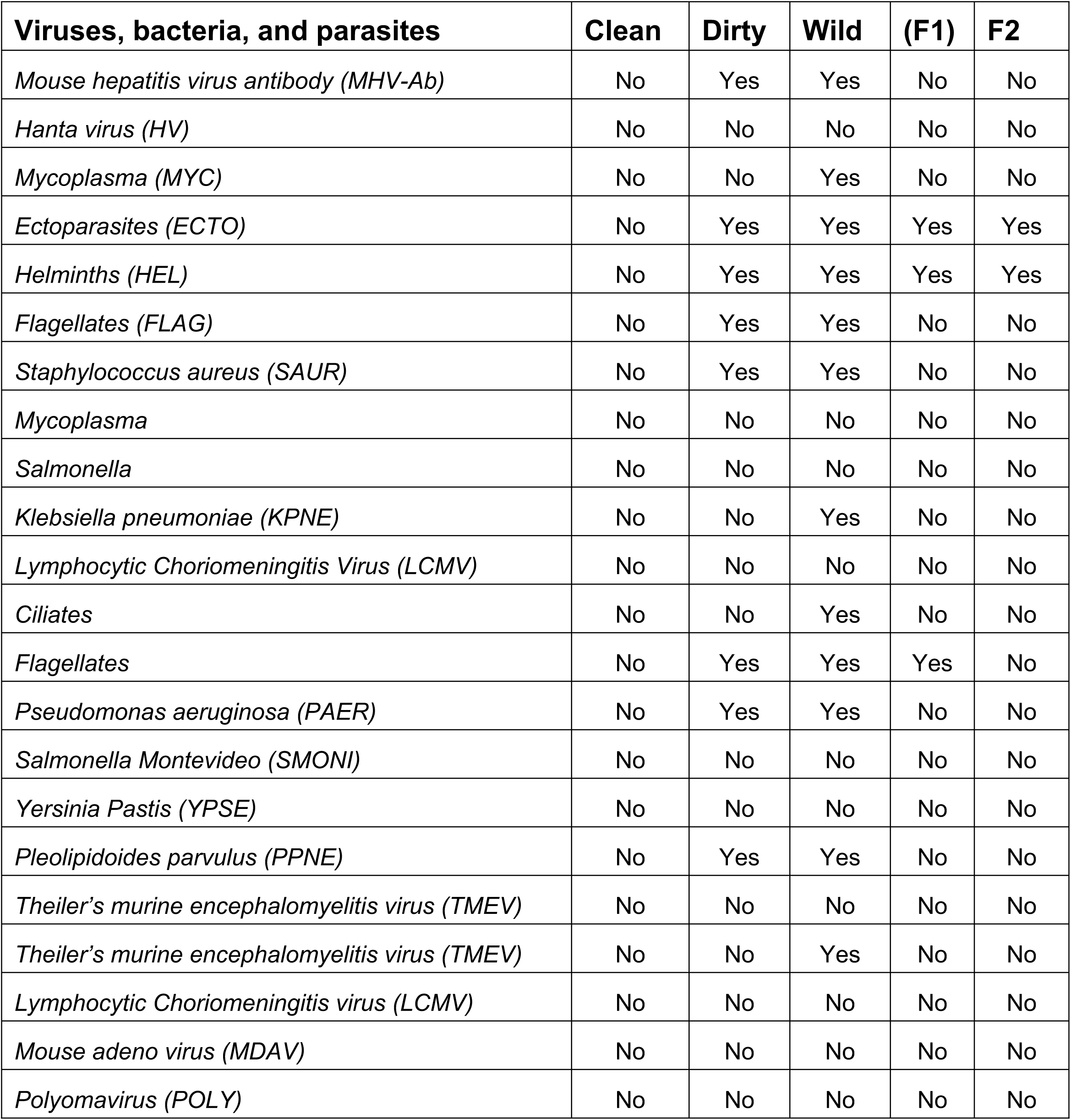
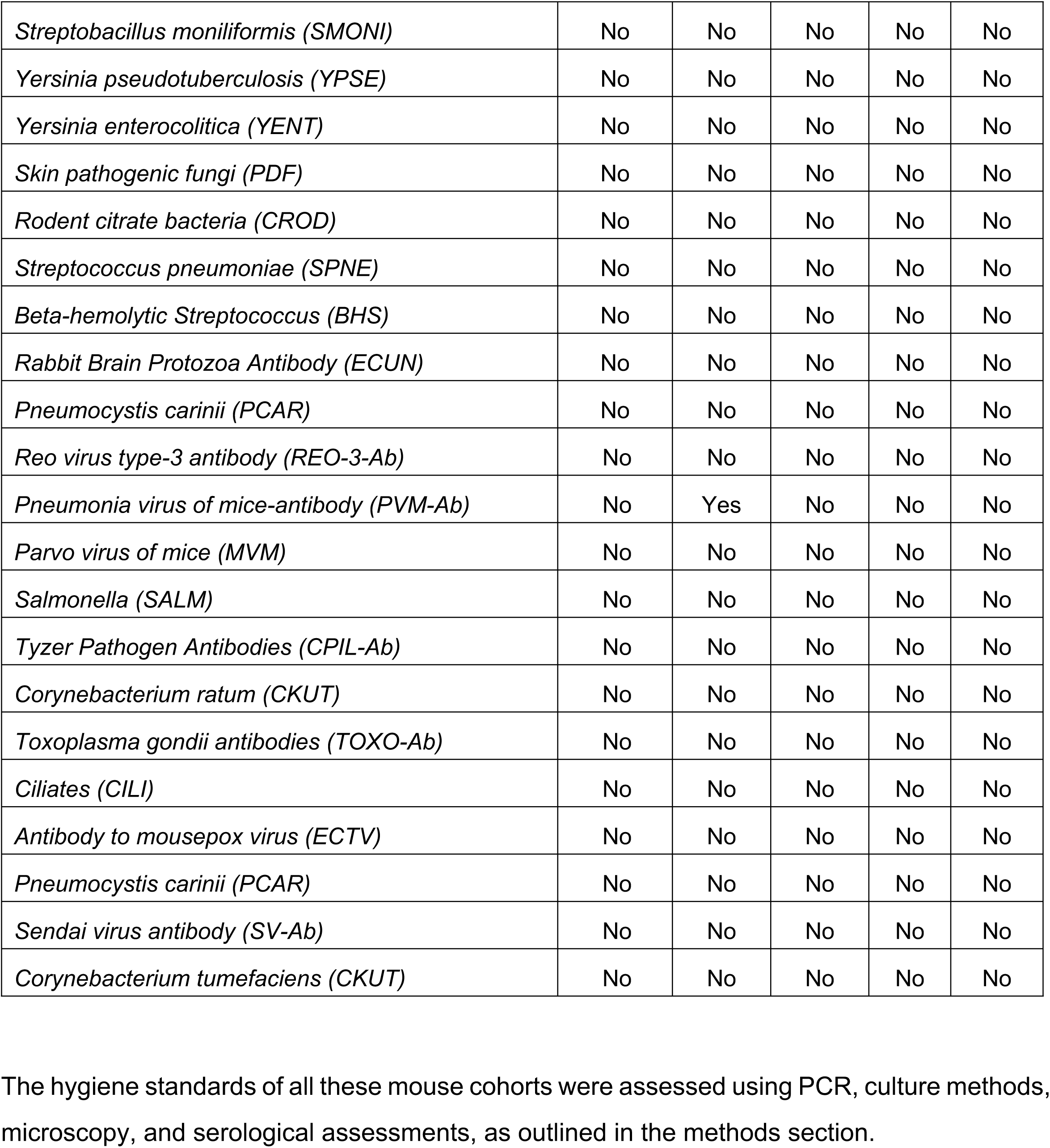
Hygiene standards of clean, dirty, field and wildlings. The hygiene standards of all these mouse cohorts were assessed using PCR, culture methods, microscopy, and serological assessments, as outlined in the methods section.

| Viruses, bacteria, and parasites | Clean | Dirty | Wild | (F1) | F2 |
| --- | --- | --- | --- | --- | --- |
| Mouse hepatitis virus antibody (MHV-Ab) | No | Yes | Yes | No | No |
| Hanta virus (HV) | No | No | No | No | No |
| Mycoplasma (MYC) | No | No | Yes | No | No |
| Ectoparasites (ECTO) | No | Yes | Yes | Yes | Yes |
| Helminths (HEL) | No | Yes | Yes | Yes | Yes |
| Flagellates (FLAG) | No | Yes | Yes | No | No |
| Staphylococcus aureus (SAUR) | No | Yes | Yes | No | No |
| Mycoplasma | No | No | No | No | No |
| Salmonella | No | No | No | No | No |
| Klebsiella pneumoniae (KPNE) | No | No | Yes | No | No |
| Lymphocytic Choriomeningitis Virus (LCMV) | No | No | No | No | No |
| Ciliates | No | No | Yes | No | No |
| Flagellates | No | Yes | Yes | Yes | No |
| Pseudomonas aeruginosa (PAER) | No | Yes | Yes | No | No |
| Salmonella Montevideo (SMONI) | No | No | No | No | No |
| Yersinia Pastis (YPSE) | No | No | No | No | No |
| Pleolipidoides parvulus (PPNE) | No | Yes | Yes | No | No |
| Theiler's murine encephalomyelitis virus (TMEV) | No | No | No | No | No |
| Theiler's murine encephalomyelitis virus (TMEV) | No | No | Yes | No | No |
| Lymphocytic Choriomeningitis virus (LCMV) | No | No | No | No | No |
| Mouse adeno virus (MDAV) | No | No | No | No | No |
| Polyomavirus (POLY) | No | No | No | No | No |
| *Streptobacillus moniliformis* (SMONI) | No | No | No | No | No |
| *Yersinia pseudotuberculosis* (YPSE) | No | No | No | No | No |
| *Yersinia enterocolitica* (YENT) | No | No | No | No | No |
| *Skin pathogenic fungi* (PDF) | No | No | No | No | No |
| *Rodent citrate bacteria* (CROD) | No | No | No | No | No |
| *Streptococcus pneumoniae* (SPNE) | No | No | No | No | No |
| *Beta-hemolytic Streptococcus* (BHS) | No | No | No | No | No |
| *Rabbit Brain Protozoa Antibody* (ECUN) | No | No | No | No | No |
| *Pneumocystis carinii* (PCAR) | No | No | No | No | No |
| *Reo virus type-3 antibody* (REO-3-Ab) | No | No | No | No | No |
| *Pneumonia virus of mice-antibody* (PVM-Ab) | No | Yes | No | No | No |
| *Parvo virus of mice* (MVM) | No | No | No | No | No |
| *Salmonella* (SALM) | No | No | No | No | No |
| *Tyzer Pathogen Antibodies* (CPIL-Ab) | No | No | No | No | No |
| *Corynebacterium ratum* (CKUT) | No | No | No | No | No |
| *Toxoplasma gondii antibodies* (TOXO-Ab) | No | No | No | No | No |
| *Ciliates* (CILI) | No | No | No | No | No |
| *Antibody to mousepox virus* (ECTV) | No | No | No | No | No |
| *Pneumocystis carinii* (PCAR) | No | No | No | No | No |
| *Sendai virus antibody* (SV-Ab) | No | No | No | No | No |
| *Corynebacterium tumefaciens* (CKUT) | No | No | No | No | No |

**Table 2.** List of antibodies for flow cytometry analysis.

| Antibodies | Source | Identifier |
| --- | --- | --- |
| Anti-mouse CD19- PE (ID3) | Biolegend | 557399 |
| Anti-mouse CD45/B220 Percp/Cy5.5 (RA3-6B2) | Biolegend | RA3-6B2 |
| Anti-mouse CD138 APC (281-2) | Biolegend | 142505 |
| Anti-mouse CD27 FITC (LG.3A10) | Biolegend | 124207 |
| Anti-mouse CD11b PE (M1/70) | Biolegend | 101207 |
| Anti-mouse CD11c APC (N418) | Biolegend | 117309 |
| Anti-mouse CDLy6G APC/Cy7 (1A8) | Biolegend | 127623 |
| Anti-mouse CDF4/80 FITC (RM8) | Biolegend | 123107 |
| Anti-mouse CDGL7- PE/Cy7 (GL7) | Biolegend | 144619 |
| Anti-mouse CD19- PE (6D3) | Biolegend | 124207 |
| Anti-mouse BV650- IgG1 (RMG1-1) | Biolegend | 406629 |
| Anti-mouse IgE- FITC (RME-1) | Biolegend | 406901 |
| Anti-mouse IgG2b- PE/cy7 (RMG2B-1) | Biolegend | 406713 |
| Anti-mouse CD8- APC/Cy7 (YTS156.7.7) | Biolegend | 126619 |
| Anti-mouse CD4- FITC (RM4-5) | Biolegend | 100509 |
| Anti-mouse CD49d- APC (R1-2) | Biolegend | 103621 |
| Anti-mouse CD44- Pacific blue (RM4-4) | Biolegend | 116008 |
| Anti-mouse Alexa 488-Foxp3 (I50D) | Biolegend | 320011 |
| Anti-mouse CD62L- PE (MEL-14) | Biolegend | 104407 |
| Anti-mouse CDICOS- APC (7E.I7G9) | Biolegend | 117419 |
| Anti-mouse CD-0X-40- PE/Cy7 (OX-86) | Biolegend | 119415 |
| Anti-mouse CD-IgG2b- PE/Cy7(RMG2b-1) | Biolegend | 406713 |

## DISCUSSION

The interaction between the immune system, environmental exposures and the gut microbiome is increasingly acknowledged as a crucial factor in health and disease [55, 56]. Laboratory mice housed under SPF conditions have been instrumental in biomedical research. However, recent studies have shown that SPF conditions lack the stimulating environment that drives the expansion and maturation of the immune system, as found in human populations [9, 10]. It has been argued that the low success rate for translating therapies from animal studies to the clinic may, in part, be due to the use of SPF mice with an immune system that, at adulthood, resembles that of a human newborn [57]. Prior studies in laboratory mice have investigated the impact of antigenic stimulants, exposure to a series of microbes, and other natural-rewilding conditions to decipher their effect on specific immune responses [31, 58, 59]. To increase the efficacy and reproducibility of mouse models in future clinical testing, mice should be exposed to more natural conditions that stimulate the immune system, thereby better mimicking the human situation. In the present study, we developed two distinct mouse models which we called “dirty” (from a facility with substandard health conditions) and “wildlings” that display different degrees of environmental and microbial exposures, and assessed the extent to which those different conditions drive adult immunity and gut microbiome composition.

Analysis of the gut microbiome revealed significant differences in the composition and diversity of bacterial species. The gut microbiota of dirty, field, and wildlings showed notable variations compared to that of clean mice. Interestingly, the microbiome composition of wildlings showed significant divergence from both field and clean mice, suggesting that this disparity could be in response to either distinct environmental variations or divergence in diets under laboratory conditions. For instance, studies have shown that not only host physiology and dietary patterns, but also environmental factors, can affect the composition of the gut microbiome [60, 61]. Environmental factors, such as temperature variations [65] and oxygen concentration [66], can also drive shifts in microbial communities in fields and wildlings. These environmental differences may contribute to the unique microbiota observed in our wildlings. This aligns with prior published studies showing that the microbiome of field mice diverges in a laboratory environment [62], in agreement with the observed divergence of microbiomes in our study.

In contrast, laboratory mice are fed on standardized diets and housed in controlled environments with regulated temperature, humidity, light cycles, and are free from predators, which limits the potential divergence of their microbiome [63]. Further serology and parasitology analysis revealed notable differences in the persistence of viruses, bacteria, and parasites at epithelial barrier sites (Table S1), indicating the significance of environmental exposure in shaping the parasite colonization. We speculate that wildlings, born from feral mothers but raised in the laboratory environment, may represent an intermediate population between field and SPF-clean mice, with adaptations that may limit pathogen colonization or suppress mechanisms that allow infections. In contrast, dirty mice live in proximity to other rodents (e.g., rats, rabbits) and were likely exposed to a more complex array of pathogens and parasites. These differences in pathogen and ectoparasite colonization between the three mouse cohorts likely reflect adaptations to conditions within distinct environments that affect the contents of microbial and parasitic exposures.

In addition to differences in gut microbiome composition, it has been appreciated in recent studies that a variety of environmental exposures and rearing conditions significantly affect the immune system [9, 21, 64, 65]. We conducted a wide range of flow cytometry analyses to investigate the effects of diverse microbial exposure and heterogeneity of the enteric gut microbiota on the differential expansion and maturation of the immune system. Our findings revealed that **‘‘**dirty**’’** mice and **‘‘**wildlings**’’** displayed varied degrees of changes in their cellular and humoral immune responses. The spleens of dirty mice exhibited a higher spleen index and displayed elevated percentages of differentiated, class-switched, plasma, memory and germinal center B cells. On the other hand, wildlings showed significantly higher proportions of mature (CD19+B220+) and differentiated B cell subsets, alongside the increased frequencies of differentiated B cell subsets. The disparity of immunological features among the three mouse cohorts suggests a strong and divergent immune response to different microbial exposures. In addition, serology assessment further suggests that high pathogen exposure (viruses, bacteria, and parasites) in substandard husbandry conditions drove the heightened activation and expansion of humoral immune response found in “dirty” mice compared to wildlings. On the other hand, our phenotypic analysis also suggests that dirty and wildlings share key immunological features, such as elevated frequencies of differentiated B cells, plasma cell abundance, and elevated GC response, even though several of these remained stronger in dirty mice compared to wildlings, suggesting similarities between the two models. Although we observed lower proportions of mature (CD19+B220+) B cells in the spleens of dirty mice, their higher expansion of B cell-specific phenotypes could be a result of the diverse array of microbial exposures, environmental stimulations, or persistence of chronic inflammation to which these mice were subjected, which drives the expansion of humoral immune responses [31]. Our findings align with recent studies by Chen and colleagues, suggesting that re-wilding conditions with natural exposure facilitate the maturation and expansion of B cells [10].

In addition to differences in B cell profiles, we observed a disparity among the frequencies of antigen-experienced T cells, which were significantly elevated in the spleens of dirty mice but relatively less in wildlings. Wildlings exhibited significantly elevated proportions of naïve T cells (CD4+CD8-, CD8+CD4-) alongside increased abundance of true memory (TMT) T cells. In contrast, dirty mice exhibited significantly elevated frequencies of all classes of antigen-experienced T cells (CD8-TTM, TvM), along with a concomitant reduction in the proportion of naïve T cells (CD8+CD4-, CD4+CD8-). One possible implication of an inflated pool of T cell populations under dirty husbandry can be attributed to their chronic and persistent exposure to various pathogens at epithelial barrier sites, leading to robust and activated T cell responses that combat these pathogens. Additionally, the diverse array of enteric microbiota in these dirty husbandry conditions may induce the secretion of short-chain fatty acids, which could in turn promote the differentiation of T cells [9]. Our studies align with those of Bura and colleagues, who previously observed activated T cell populations in pet-store mice [20], in line with the impact of an altered environment on the expansion of these cells [65]. Although less exposed than dirty mice, wildlings also exhibited elevated frequencies of true memory T cells (CD8+CD44+CD49d+), suggesting the establishment of specific memories from prior exposure to infections [66], which were preserved even after housing in an environment with fewer germs than that of dirty mice. Similar studies conducted by Yeung and colleagues reported that laboratory mice subjected to enriched natural exposure displayed elevated proportions of different antigen-experienced T cells [65], underscoring the notion that gut microbiota composition and enriched natural exposure can drive the expansion of TMT cells. In addition, we observed increased frequencies of naïve T cells in wildlings, suggesting the existence of differential mechanisms contributing to the expansion of these cell types in this cohort. In a recent study, Ma and colleagues also reported increased frequencies of naïve T cells in wildlings, although, different from our study, a significant elevation in the proportions of antigen-experienced T cells was also observed [21]. This divergence could be attributed to the age of the mice analyzed, variations in laboratory husbandry conditions, early-life exposure to natural supplements (such as soil and hay), or the distinct microbial and pathogenic exposures of the founding feral populations.

With regards to myeloid cells, while we observed a significant expansion in dirty mice, no significant differences were found between lab mice and wildlings, highlighting a significant difference between dirty and wilding model systems. One possible reason is that wildlings, while exposed to ectoparasites, may not have encountered the same magnitude, variety and persistence of pathogen exposure as dirty mice, which would more readily activate myeloid cells. It is also possible that laboratory husbandry conditions may have induced wildlings to prioritize adaptive immune responses without triggering widespread systemic activation of myeloid cells. As endogenous expansion of CD11b+ myeloid cells has been reported in inflammatory situations to fight infections [67], it is likely that chronic exposure to pathogens in the substandard housing environment to which dirty mice were exposed induced the secretion of endogenous cytokines, such as IL-17 and granulocyte colony-stimulating factor (G-CSF), that increase granulopoiesis [68, 69] in the these mice. In contrast, wildlings were only colonised by ectoparasites, and they were not continuously exposed as adults in the same manner as dirty mice.

Lastly, we observed a decrease in the colonisation of ectoparasites in F2 progenies of wildlings (Table:1). We further detected a reduction in specific immune cell subsets such as activated B cells, as well as CD8-TTM and CD8-TvM in F2 progenies of wildlings. These observations suggest that the lack of a continued stimulating environment (i.e., pathogens at epithelial and barrier sites, stressful housing conditions, etc.) can lead to reduced levels of these cells as the immune system adapts to a less challenging environment. Similar studies have previously shown how diet or stress conditions affect transgenerational immune phenotypes [70]. Thus, it is likely that either the modification in diet, microbiome, or the absence of chronic stressors and continued microbial exposure resulted in a diminished presence of activated immune cell signatures in F2 progenies. For the wildling model of immune system activation to be useful, future studies should focus on developing conditions that enable reproducible transgenerational transfer and maintenance of active immune features in wildling colonies.

In summary, our findings underscore the impact of diverse microbial exposures and other environmental conditions on adult immunity. Wildlings and different approaches (such as pet stores, rewilding, feralization, and cohousing) that expose laboratory mice to environmental variables will be particularly useful for investigating immune responses and their broader impact on animal physiology. However, we caution that, as highlighted in our study, intrinsic variations in microbial exposure can influence the composition of innate and adaptive immune responses and thus affect the reproducibility of studies performed at different laboratories using different model systems. Identifying the specific effects of distinct exposures and housing conditions on the expansion and maturation of innate and adaptive immune features, although challenging, will be a necessary objective of future studies if mouse cohorts with immune systems activated beyond SPF levels are to become reliable and useful models of human disease and for research in a variety of areas such as metabolism, cancer and neurodegeneration.

## Material and methods

### “Clean”, ‘‘dirty’’ and “wildling” mouse models

3-week-old C57BL/6 standard laboratory mice (clean) were acquired from Charles River and bred on-site under SPF (specific pathogen-free) conditions, free from murine norovirus (MNV) and Helicobacter, at the Chinese Institute for Brain Research (CIBR), Beijing, adhering to strict characterization SPF standards. Mice were kept under a regular and controlled 12-h day/night cycle with ad libitum access to food and water. Mice bred under SPF conditions are herein referred to as “clean”. Mice herein referred to as “dirty” were acquired from an animal facility in Southern Beijing which holds several other rodents (e.g., rats, rabbits) in the same quarters and under substandard hygienic conditions. Animals herein referred to as “wildings” are C57Bl/6 mice born after embryo transfer from females captured in the field using inverse germ-free rederivation techniques as outlined by Rosshart and colleagues *[24]*. To this end, wild mice (*M. musculus domesticus*) were captured and trapped from geographically distinct regions (Shandong and Henan provinces) of China using live animal aluminium traps as described previously [71]. The mouse-catching traps were monitored regularly to prevent prolonged captivity and the trapping of immature mice [71].

Typically, wild mice were pre-selected and screened based on their size and color appearance to exclude rats and other species, and 30-40% were sent to commercial veterinary research laboratories (VRL) in Suzhou, China, for health assessment, including analysis of ectoparasites, bacteria, viruses, fungi, and other commensal pathogens at the epithelial and barrier sites. The remaining mice were sent to a non-SPF site (i.e., ABSL2) of CIBR’s animal facility and divided into separate cages. To generate wildling mice, C57BL/6 embryos from the SPF side of the facility were frozen in liquid nitrogen cylinders for later surgical transfers [24]. Pseudopregnant field female mice were generated by crossing with surgically vasectomized male field mice. Subsequently, SPF mouse embryos were surgically implanted into several batches of pseudopregnant field mice. More than 30% of these mice gave birth to healthy offspring, herein referred to as wildlings. Beddings from field mice were seeded in the cages of weaned wildlings to further increase their microbial exposure during adolescence. Wildlings and dirty mice were self-crossed to obtain F2 wildling progenies. Wildling mice were housed in the ABSL2 site of CIBR’s animal facility and fed with carrots, soybeans, corn and other components of their natural diet. Three wildlings were raised in one cage after weaning to reduce the likelihood of them fighting with each other. Three-month-old male mice from the clean, dirty and wilding cohorts were used in this study. All animal experiments were conducted in accordance with the protocol approved by CIBR’s IACUC (CIBR-IACUC-061).

### Serology and parasitology analyses

Randomly selected mice from clean, dirty, field and wildlings were subjected to a health assessment for the presence of infectious agents and parasites using PCR, ELISA, and microbial cultures. Faecal samples were desiccated on Opti-Spot cards, following the manufacturer’s sampling protocols (IDEXX BioAnalytics). Oral and other bodily swabs were obtained according to the sample submission protocols of Charles River Laboratories. Whole blood was obtained from the tail vein of mice and placed into serum separator tubes. Samples were left to coagulate (30–60 minutes, ambient temperature) and subsequently centrifuged (10 minutes, 1,500–2,000 × g). Serum was distributed into DNase/RNase-free tubes to prevent haemolysis and was preserved at −80 °C until dispatch. Batched aliquots (≥100 µL per animal) were dispatched to a commercial diagnostic veterinary research laboratory (VRL, Suzhou, China) on dry ice. Mouse blood serology was performed by ELISA utilising manufacturer-validated kits. The test panel comprised prevalent murine infections, including mouse hepatitis virus (MHV), mouse norovirus, parvoviruses, Sendai virus, pneumonia virus of mice, Mycoplasma pulmonis, Theiler’s encephalomyelitis virus (TEV), ectromelia, and EDIM/rotavirus, as specified in the VRL request. Details of the pathogens and parasites screened can be found in Table 1. Samples exhibiting significant haemolysis or inadequate volume were eliminated. The experiments were conducted on pooled samples; a microorganism was deemed present if recognized by at least one method.

### 16S rRNA microbiome sequencing

Gut microbiome analysis was performed as previously described (Wang et al., 2025, BMC Biology PMID: 40346662). Briefly, fecal DNA was extracted using a commercial kit (TianGen, China, Catalog #: DP336) and the hypervariable V4 region of the 16S rRNA gene was amplified with barcoded 515F/806R primers. PCR was performed using Phusion® High-Fidelity Master Mix (New England Biolabs). Amplified products were verified via electrophoresis, pooled in equidensity ratios, and purified with the Universal DNA Purification Kit (TianGen, China, Catalogue #: DP214). Sequencing libraries were prepared using the NEBNext Ultra II DNA Library Prep Kit (Cat. No. E7645B, New England Biolabs) and validated by Qubit (ThermoFisher Scientific, USA) and qPCR before being sequenced on an Illumina NovaSeq 6000 system (Illumina Novaseq6000, Illumina, San Diego, CA, USA). Bioinformatic processing was conducted in Quantitative Microbial Ecology (QIIME2, version 2022.02) [7].

### Flow cytometry

Peripheral blood was collected from the tail of mice and kept in an EDTA-containing tube on ice. For red blood cell lysis, 300 μL of blood and spleen suspension were resuspended in red blood cell lysis buffer (BD Pharm LyseT^m^ Cat: 555899) for 10 minutes. The whole spleen and mesenteric lymph node (MLN) were isolated from their surrounding abdominal area and cleaned with a surgical scalpel and paper towels to remove surrounding hairs and blood contamination in 1x PBS. Spleens were immersed in ice-cold RPMI-1640 media (Gibco) supplemented with 2% foetal bovine serum (FBS; Gibco) and 1% penicillin-streptomycin (Gibco). Each spleen and MLN were placed in a 70-µm cell strainer (BD Falcon) positioned above a 50-mL tube containing 5–10 mL of FACS buffer. Tissues were mechanically separated by gently compressing with the plunger end of a sterile 5-mL syringe until single-cell suspensions were achieved. The suspensions were subsequently filtered through a 70μm strainer (Falcon) and washed with an additional 5 mL of medium to optimize cell recovery. To reduce non-specific antibody binding, cells were initially treated with anti-mouse FcR blocking reagent (Fc block; Lot: 5241207109, Miltenyi) for 30 minutes at 4 °C. Next, single-cell suspensions were counted using a cell counter (automatic) after staining with trypan blue (Thermo Fisher Scientific). Cell viability was determined using 7-AAD (Biolegend: Catt:420404) and DAPI (Solarbio: Catt: C0065).

The cell suspensions were preincubated with the respective cell surface-specific antibodies in stain buffer (BD Pharmingen, Cat: 554657) at 4°C for 20 minutes, followed by incubation at room temperature to stain the cell surface markers. A detailed list of all cell surface and intracellular markers analyzed is provided in (Table S2). To intracellularly stain the IgG1, IgG2B, Intracellular staining with Brilliant™ Violet (BV-650) conjugated IgG1 (Biolegend, Cat:406629), and Phycoerythrin (PE/Cy7)-conjugated anti-mouse IgG2b (Biolegend, Cat:406713) antibodies was performed using the Cytofix/Cytoperm fixation/permeabilization kit (BD Pharmingen, Cat: 554714) following the manufacturer’s instructions. Analytical flow cytometry was performed on the BD LSR Fortessa (BD Biosciences). Controls included single-stained and negative samples. Fluorescence-minus-one (FMO) controls were used to establish gating thresholds. Each sample recorded a minimum of 100,000 live, single events. Flow cytometry data were analysed utilising FlowJo software (version 10.8.1, BD Biosciences). Populations were delineated using sequential gating based on forward/side scatter, singlet discrimination (FSC-A against FSC-H), exclusion of necrotic cells, and lineage-specific markers. The frequencies of B-cell, T-cell, and myeloid subgroups were represented as percentages of total viable splenocytes or relevant parent populations.

### Statistical analysis

The statistical analysis employed is specified in each figure and was performed using the GraphPad Prism 9.5.1 software. To determine the statistical significance, we used a one-way non-parametric ANOVA. We considered differences statistically significant when p < 0.05. We indicate significance levels with asterisks: * for p< 0.05, ** for p< 0.01, *** for p< 0.001, and **** for p< 0.0001. All plots show the standard error of the mean (SEM).

## Acknowledgments

The authors would like to thank Lei Wang, Jocelyn Jia, Yankui Fu and Shuo Zhang for technical and admin assistance.

## Ethics statement

All animal experiments were performed in compliance with the protocol approved by the Institutional Animal Care and Use Committee (IACUC) of the Chinese Institute of Brain Research (CIBR-IACUC-061).

## Authors’ contributions

KH: performed all experimental work, analyzed data and prepared a draft of the manuscript and figures. MX: co-supervised the project and corrected the manuscript. CFI: supervised the project and corrected the manuscript. This work was supported by research grants to C.F.I. from Peking University, Chinese Institute for Brain Research, Beijing, and Swedish Research Council (Vetenskapsrådet, contract nr. 2024-03222); and a startup grant to M.X. from Swedish Research Council (Vetenskapsrådet, contract nr. 2021-01805).

## Data availability statement

The data that support this work are available from the corresponding author upon request.

## Supplementary Figure legend and tables

**Figure S1.**
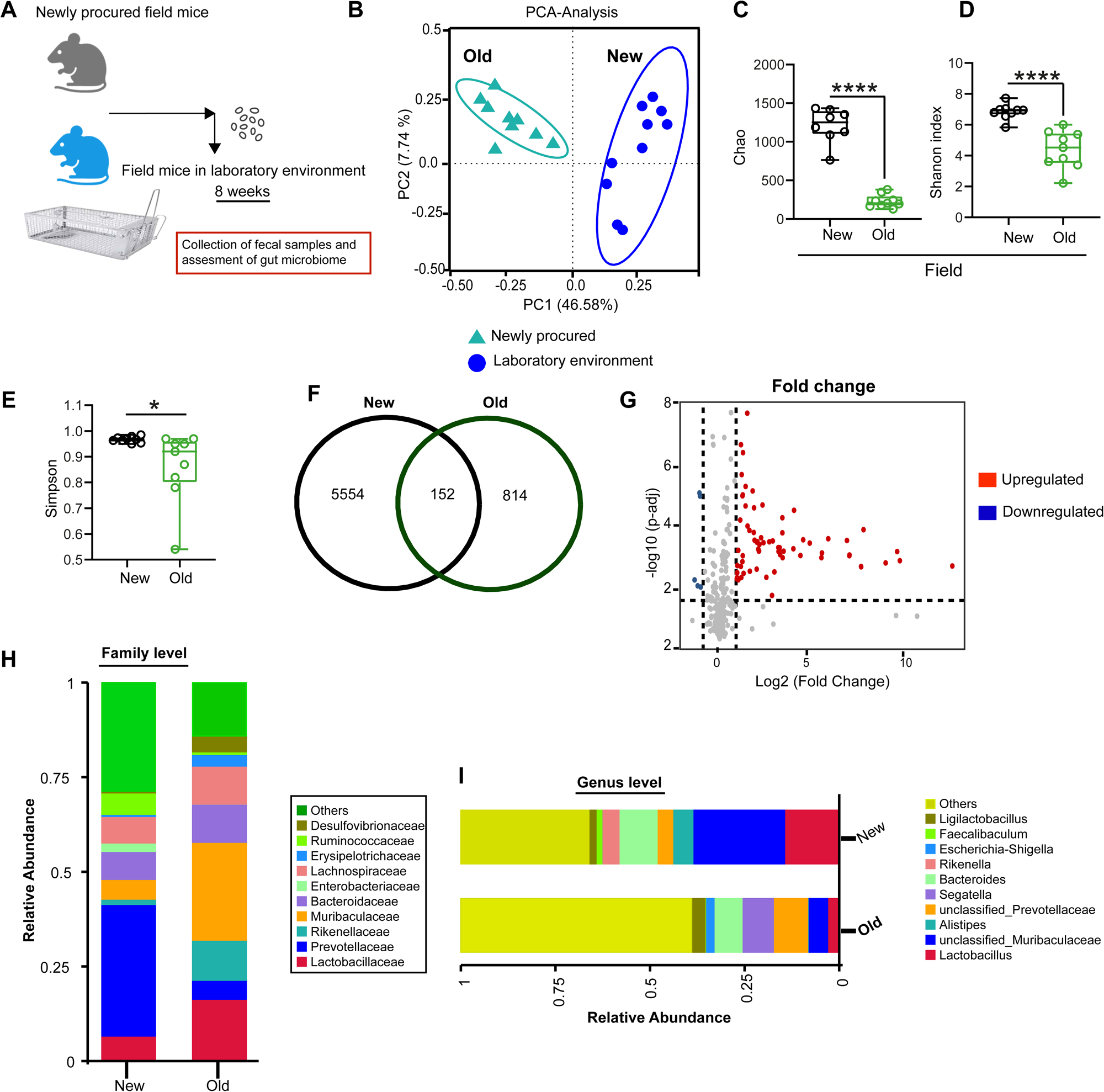
Altered composition of the gut microbiome of field mice after laboratory housing (supplement to Figure 1) **(A)** Schematic figure showing comparative assessment of gut microbiome of newly procured field mice and field mice after prolonged housing in the non-SPF laboratory environment. **(B)** The principal component analysis (PCA) plot illustrates the beta-diversity of the gut microbiomes of field mice (new vs. old). **(C-E)** Statistical bar plot represents alpha diversity metrics, Chao index **(C)**, Shannon index **(D)**, and Simpson index **(E)** among the gut microbiomes of both Field cohorts. **(F)** Venn diagram shows the shared and different species among the gut microbiome of both field mouse groups. **(G)** Fold change graphs show the significantly upregulated and downregulated microbiome species in both field mouse groups. **(H, I)** Relative abundance of bacteria at the family level (**H**) and genus **(I)** levels in gut microbiome of both mouse cohorts. Dots in bar graphs correspond to individual mice. Means are shown. Unpaired two-tailed t-tests determined statistical significance and P values.

**Fig. S2.**
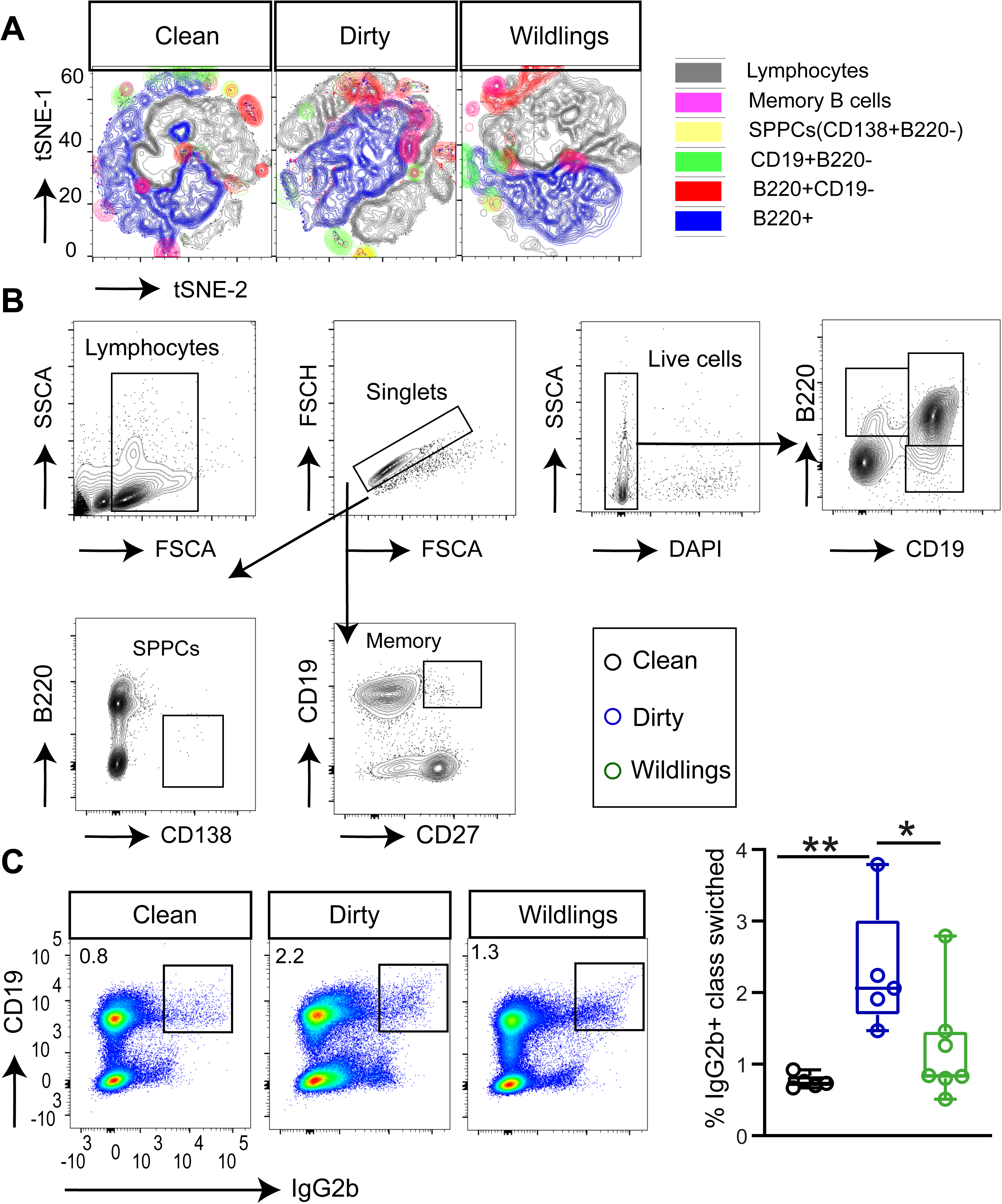
Distinct effects of altered microbial exposures and housing conditions on the expansion of B cells (supplement to Figure 2) **(A)** Different B cell subsets were quantified by flow cytometry based on the gating strategy described from clean, dirty, and wildling mice and visualized with t-SNE showing the distribution of different naïve and antigen-experienced B cell subsets. **(B)** Representative flow cytometry contour plots depicting the gating strategy for B220+CD19-, CD19+B220-, memory (CD19+CD27+), SPPCs (CD138+B220-) and B220+ B cells in the spleen from clean mice. Each colour represents B cell subset. **(C)** Representative flow cytometry pseudocolour plots depicting the CD19 vs. IgG2b expression in gated DAPI negative live cells (left), and summary statistic plot (right) shows the frequencies of CD19+IgG2b+ class-switched cells in the spleens of clean, dirty, and wildling mice. Each dot represents a single mouse. Data are expressed as mean SEM. Statistical quantification among different groups was analyzed by non-parametric one-way ANOVA with Tukey’s multiple comparison tests; *p < 0.05; **p < 0.01.

**Fig. S3.**
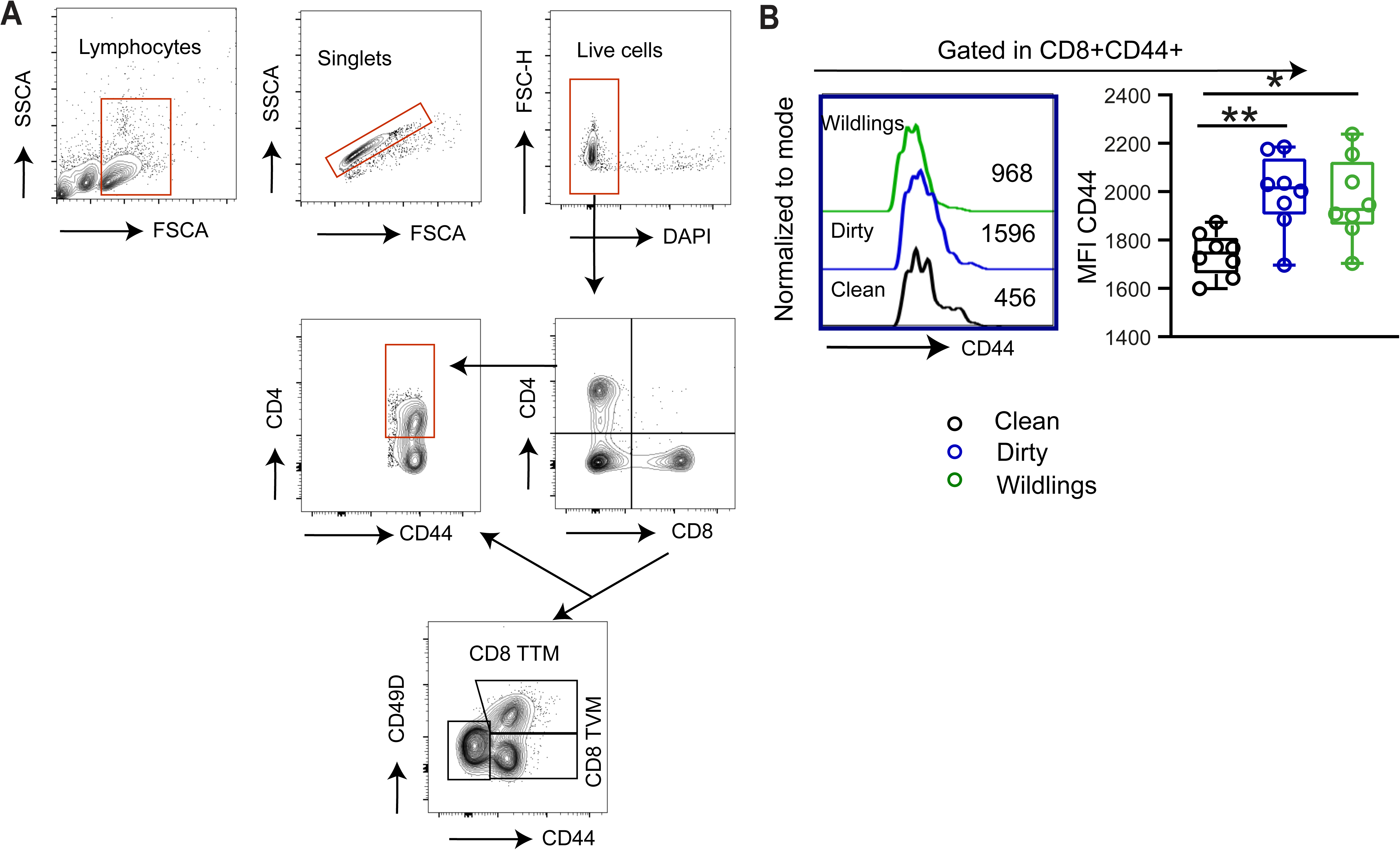
Effects of altered microbial exposures and housing conditions on spleen T cell pools (supplement to Figure 3) **(A)** Representative contour flow cytometry plots depicting the gating strategy to characterize different naïve (CD4+CD8-, CD8+CD4-) and antigen-experienced T cells (TTM, TVM) in the spleen of clean, dirty and wildlings. **(B)** Histogram (middle) depicts the CD44 expression and cell counts gated in CD8+CD44+ cells, and bar graph (right) shows quantification of the mean fluorescence intensity (MFI) of CD44 values in the spleen of clean, dirty and wildlings. Each dot represents a single mouse. Data are expressed as mean SEM. Statistical differences among the different groups were analyzed using non-parametric one-way ANOVA (Tukey’s multiple comparison tests); *p < 0.05, **p < 0.01.

**Fig. S4.**
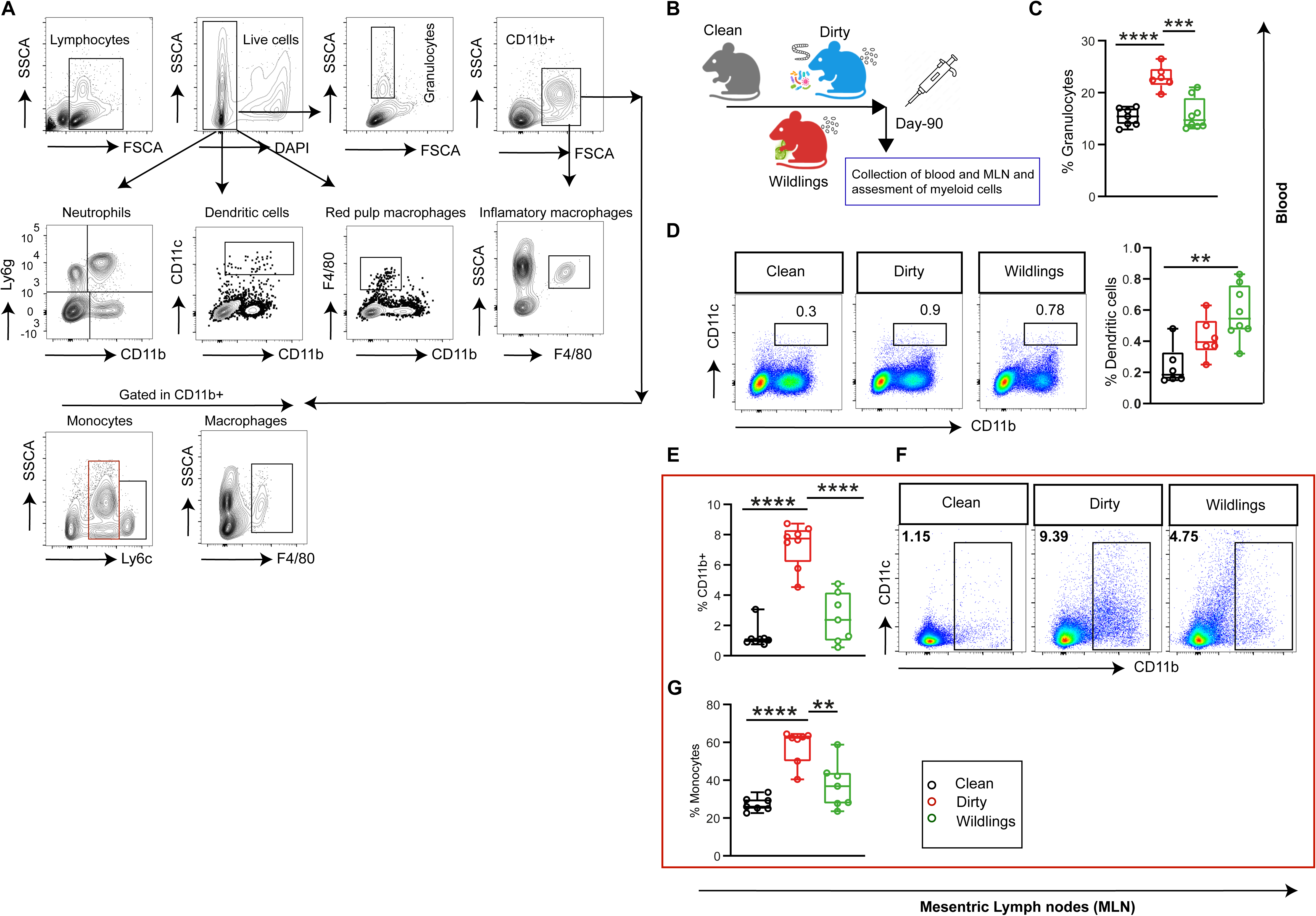
Effects of environmental conditions on myeloid cells (supplement to Figure 4) **(A)** Representative flow cytometry plots illustrate the gating strategy used to characterize different myeloid cell subsets in the spleen. **(B)** Schematic overview of clean, dirty and wildlings. Peripheral blood was collected at postnatal day (p-90) to profile different myeloid cell subsets. **(C)** Frequencies of granulocytes (FSC^high^SSC^high^) in the blood of all mouse cohorts. **(D)** Representative pseudocolour flow cytometry plots (left) depict the CD11b vs CD11c expression (left) and summary statistic plots (right) present the frequencies of dendritic cells (CD11C^high^, CD11b^high^) in the blood of clean, dirty and wildlings. **(E)** Graphs depict the frequencies of CD11b+ myeloid cells in the MLN of clean, dirty and wildling cohorts. **(F, G)** Representative flow cytometry pseudocolour plots illustrate CD11b expression **(F)** and summary statistic plot shows the quantification of the frequencies of monocytes (Ly6c^high^Ly6c^low^CD11b+) cells **(G)** in the MLN of clean, dirty and wildling cohorts. Each dot represents a single mouse. Data are expressed as mean SEM. Statistical quantification among different groups was analyzed using non-parametric ANOVA (Tukey’s multiple comparison tests); *p < 0.05, **p < 0.01.

**Fig. S5.**
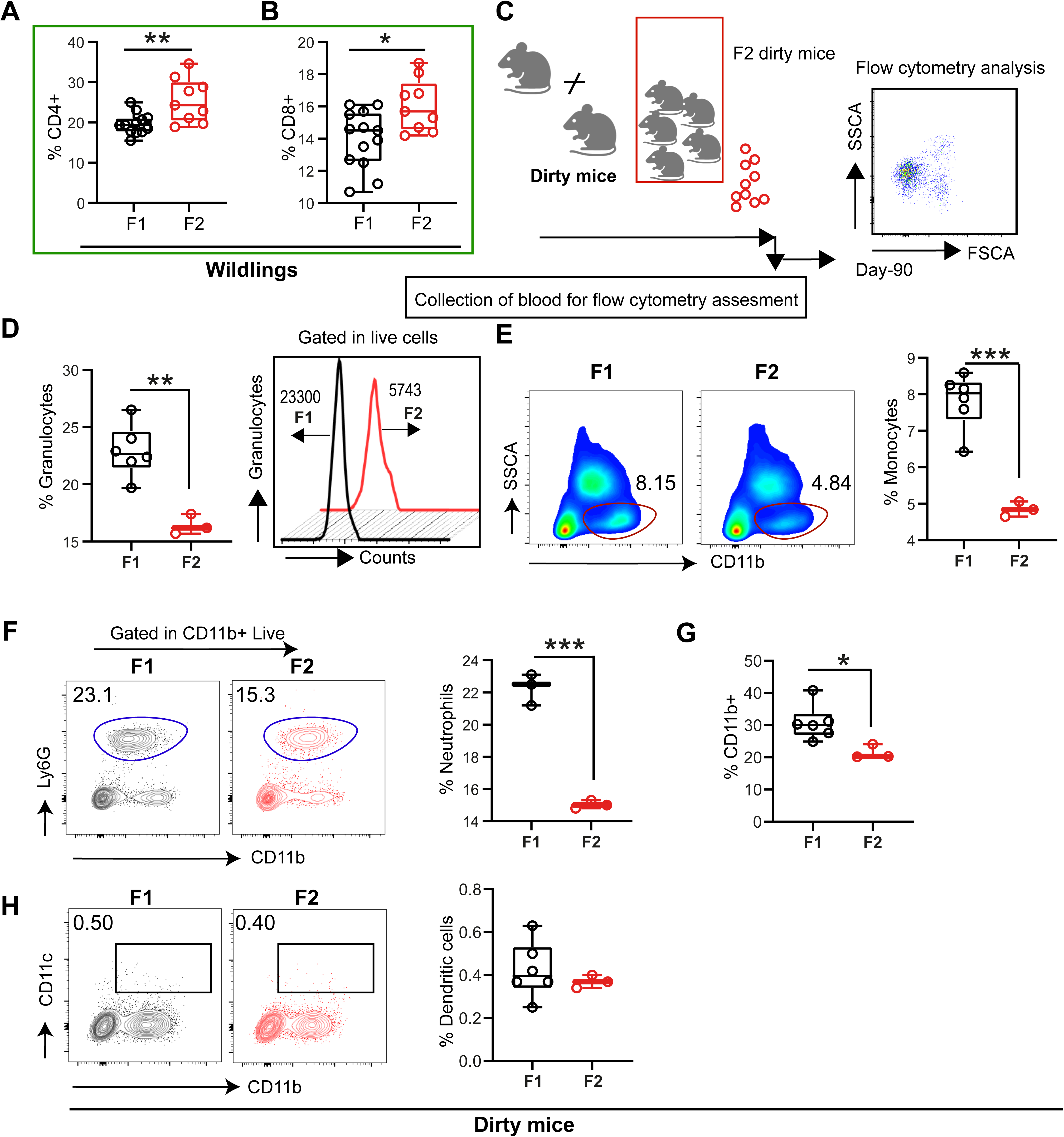
The immune phenotypes of wildlings and dirty mice converge to those of clean mice after two generations in a laboratory environment (supplement to Figure 5) **(A, B)** Summary statistic plots depict the frequencies of CD4+ **(A)** and CD8+ **(B)** T cells in the spleen of F1 and F2 generations of wildlings. **(C)** Schematic of F1 and F2 dirty mice and comparative assessment of immune phenotypes at postnatal day 90 from the peripheral circulation **(D**). Summary statistics of the frequencies of granulocytes (SSChighFSChigh) (left) and staggered histogram (middle) display the cell counts in the blood of F1 and F2 dirty mice. **(E)** Representative flow cytometry contour plots show the expression of CD11b and corresponding frequencies of monocytes in the blood of F1 and F2 dirty mice. (**F)** Representative flow cytometry plots depicting the Ly6G vs. CD11b expression (left) and corresponding frequencies of neutrophils (CD11b+Ly6G+) in the blood of F1 and F2 dirty mice. **(G)** Summary statistics of the frequencies of CD11b+ myeloid cells in the blood of F1 and F2 dirty mice. **(H)** Representative flow cytometry plots illustrate the expression of CD11c vs CD11b in the blood of F1 and F2 dirty mice (left) and summary statistic plot (right) shows the frequencies of circulating dendritic cells (CD11c^high^+CD11b^high^+). Each dot represents a single mouse. Data are expressed as mean SEM. Statistical differences among the different groups were analyzed using unpaired two-tailed t-tests; *p < 0.05, **p < 0.01.

## References

1. Graham, A.L., Naturalizing mouse models for immunology. Nature Immunology, 2021. 22(2): p. 111–117.

2. Abolins, S., et al., The comparative immunology of wild and laboratory mice, Mus musculus domesticus. Nature communications, 2017. 8(1): p. 14811.

3. Masopust, D., C.P. Sivula, and S.C. Jameson, Of mice, dirty mice, and men: using mice to understand human immunology. The Journal of Immunology, 2017. 199(2): p. 383–388.

4. Tao, L. and T.A. Reese, Making mouse models that reflect human immune responses. Trends in immunology, 2017. 38(3): p. 181–193.

5. Mestas, J. and C.C. Hughes, Of mice and not men: differences between mouse and human immunology. The Journal of Immunology, 2004. 172(5): p. 2731–2738.

6. Huggins, M.A., S.C. Jameson, and S.E. Hamilton, Embracing microbial exposure in mouse research. Journal of leukocyte biology, 2019. 105(1): p. 73–79.

7. Shay, T., et al., Conservation and divergence in the transcriptional programs of the human and mouse immune systems. Proceedings of the National Academy of Sciences, 2013. 110(8): p. 2946–2951.

8. Payne, K.J. and G.M. Crooks, Immune-cell lineage commitment: translation from mice to humans. Immunity, 2007. 26(6): p. 674–677.

9. Beura, L.K., et al., Normalizing the environment recapitulates adult human immune traits in laboratory mice. Nature, 2016. 532(7600): p. 512–516.

10. Chen, Y.-H., et al., Rewilding catalyzes maturation of the humoral immune system. Science Advances, 2025. 11(10): p. eads2364.

11. Ronan, V., R. Yeasin, and E.C. Claud, Childhood development and the microbiome— the intestinal microbiota in maintenance of health and development of disease during childhood development. Gastroenterology, 2021. 160(2): p. 495–506.

12. Perlroth, N.H. and C.W.C. Branco, Current knowledge of environmental exposure in children during the sensitive developmental periods☆. Jornal de Pediatria, 2017. 93: p. 17–27.

13. Bengmark, S., Gut microbiota, immune development and function. Pharmacological research, 2013. 69(1): p. 87–113.

14. Round, J.L. and S.K. Mazmanian, The gut microbiota shapes intestinal immune responses during health and disease. Nature reviews immunology, 2009. 9(5): p. 313–323.

15. Tremaroli, V. and F. Bäckhed, Functional interactions between the gut microbiota and host metabolism. Nature, 2012. 489(7415): p. 242–249.

16. Kamada, N., et al., Control of pathogens and pathobionts by the gut microbiota. Nature immunology, 2013. 14(7): p. 685–690.

17. Rio, P., et al., Pollutants, microbiota and immune system: frenemies within the gut. Frontiers in Public Health, 2024. 12: p. 1285186.

18. Kharrazian, D., Exposure to environmental toxins and autoimmune conditions. Integrative Medicine: A Clinician’s Journal, 2021. 20(2): p. 20.

19. Husso, A., et al., Impacts of maternal microbiota and microbial metabolites on fetal intestine, brain, and placenta. BMC biology, 2023. 21(1): p. 207.

20. Macpherson, A.J., M.G. De Agüero, and S.C. Ganal-Vonarburg, How nutrition and the maternal microbiota shape the neonatal immune system. Nature Reviews Immunology, 2017. 17(8): p. 508–517.

21. Ma, J., et al., Laboratory mice with a wild microbiota generate strong allergic immune responses. Science immunology, 2023. 8(87): p. eadf7702.

22. Pritchett, K.R., Helminth parasites of laboratory mice, in The Mouse in Biomedical Research. 2007, Elsevier. p. 551–564.

23. Baker, D.G., Natural pathogens of laboratory mice, rats, and rabbits and their effects on research. Clinical microbiology reviews, 1998. 11(2): p. 231–266.

24. Rosshart, S.P., et al., Laboratory mice born to wild mice have natural microbiota and model human immune responses. Science, 2019. 365(6452): p. eaaw4361.

25. Gollwitzer, E.S. and B.J. Marsland, Impact of early-life exposures on immune maturation and susceptibility to disease. Trends in immunology, 2015. 36(11): p. 684–696.

26. Renz, H. and C. Skevaki, Early life microbial exposures and allergy risks: opportunities for prevention. Nature Reviews Immunology, 2021. 21(3): p. 177–191.

27. Goenka, A. and T.R. Kollmann, Development of immunity in early life. Journal of Infection, 2015. 71: p. S112–S120.

28. Nielsen, C.C., et al., Natural environments in the urban context and gut microbiota in infants. Environment international, 2020. 142: p. 105881.

29. Shin, J.-H., et al., Lifestyle and geographic insights into the distinct gut microbiota in elderly women from two different geographic locations. Journal of physiological anthropology, 2016. 35(1): p. 31.

30. Rook, G.A., Regulation of the immune system by biodiversity from the natural environment: an ecosystem service essential to health. Proceedings of the National Academy of Sciences, 2013. 110(46): p. 18360–18367.

31. Suzuki, T., et al., Environmental pollutants and the immune response. Nature Immunology, 2020. 21(12): p. 1486–1495.

32. Zhang, Q., et al., Genetic mapping of microbial and host traits reveals production of immunomodulatory lipids by Akkermansia muciniphila in the murine gut. Nature microbiology, 2023. 8(3): p. 424–440.

33. Burger, S., et al., Natural microbial exposure from the earliest natural time point enhances immune development by expanding immune cell progenitors and mature immune cells. The Journal of Immunology, 2023. 210(11): p. 1740–1751.

34. Zheng, D., T. Liwinski, and E. Elinav, Interaction between microbiota and immunity in health and disease. Cell research, 2020. 30(6): p. 492–506.

35. Tellier, J. and S.L. Nutt, Standing out from the crowd: How to identify plasma cells. European journal of immunology, 2017. 47(8): p. 1276–1279.

36. Khodadadi, L., et al., The maintenance of memory plasma cells. Frontiers in immunology, 2019. 10: p. 721.

37. Shi, W., et al., Transcriptional profiling of mouse B cell terminal differentiation defines a signature for antibody-secreting plasma cells. Nature immunology, 2015. 16(6): p. 663–673.

38. Wang, K., G. Wei, and D. Liu, CD19: a biomarker for B cell development, lymphoma diagnosis and therapy. Experimental hematology & oncology, 2012. 1: p. 1–7.

39. Kurosaki, T., K. Kometani, and W. Ise, Memory B cells. Nature Reviews Immunology, 2015. 15(3): p. 149–159.

40. Cyster, J.G. and C.D. Allen, B cell responses: cell interaction dynamics and decisions. Cell, 2019. 177(3): p. 524–540.

41. Syeda, M.Z., et al., B cell memory: from generation to reactivation: a multipronged defense wall against pathogens. Cell Death Discovery, 2024. 10(1): p. 117.

42. Naito, Y., et al., Germinal center marker GL7 probes activation-dependent repression of N-glycolylneuraminic acid, a sialic acid species involved in the negative modulation of B-cell activation. Molecular and cellular biology, 2007. 27(8): p. 3008–3022.

43. Sun, L., et al., T cells in health and disease. Signal transduction and targeted therapy, 2023. 8(1): p. 235.

44. Chopp, L., et al., From thymus to tissues and tumors: A review of T-cell biology. Journal of Allergy and Clinical Immunology, 2023. 151(1): p. 81–97.

45. Wu, T.D., et al., Peripheral T cell expansion predicts tumour infiltration and clinical response. Nature, 2020. 579(7798): p. 274–278.

46. Fang, H., et al., Gut-Spleen axis: microbiota via vascular and immune pathways improve busulfan-induced spleen disruption. MSphere, 2023. 8(1): p. e00581–22.

47. Petersone, L., et al., T cell/B cell collaboration and autoimmunity: an intimate relationship. Frontiers in immunology, 2018. 9: p. 1941.

48. Kawamoto, H. and N. Minato, Myeloid cells. The international journal of biochemistry & cell biology, 2004. 36(8): p. 1374–1379.

49. Khan, S.Q., I. Khan, and V. Gupta, CD11b activity modulates pathogenesis of lupus nephritis. Frontiers in medicine, 2018. 5: p. 52.

50. Izumi, G., et al., CD11b+ lung dendritic cells at different stages of maturation induce Th17 or Th2 differentiation. Nature communications, 2021. 12(1): p. 5029.

51. Liu, Z., et al., Analysis of myeloid cells in mouse tissues with flow cytometry. STAR protocols, 2020. 1(1): p. 100029.

52. Wastyk, H.C., et al., Gut-microbiota-targeted diets modulate human immune status. Cell, 2021. 184(16): p. 4137–4153. e14.

53. Baaten, B.J., et al., Regulation of antigen-experienced T cells: lessons from the quintessential memory marker CD44. Frontiers in immunology, 2012. 3: p. 23.

54. Mrass, P., et al., CD44 mediates successful interstitial navigation by killer T cells and enables efficient antitumor immunity. Immunity, 2008. 29(6): p. 971–985.

55. Hooper, L.V., D.R. Littman, and A.J. Macpherson, Interactions between the microbiota and the immune system. science, 2012. 336(6086): p. 1268–1273.

56. Cerf-Bensussan, N. and V. Gaboriau-Routhiau, The immune system and the gut microbiota: friends or foes? Nature Reviews Immunology, 2010. 10(10): p. 735–744.

57. Willyard, C., Squeaky clean mice could be ruining research. Nature, 2018. 556(7700): p. 16–19.

58. Reese, T.A., et al., Sequential infection with common pathogens promotes human-like immune gene expression and altered vaccine response. Cell host & microbe, 2016. 19(5): p. 713–719.

59. Yan, A.W., et al., Sequential infection experiments for quantifying innate and adaptive immunity during influenza infection. PLoS computational biology, 2019. 15(1): p. e1006568.

60. Neha, S.A., et al., Impacts of host phylogeny, diet, and geography on the gut microbiome of rodents. Plos one, 2025. 20(1): p. e0316101.

61. Goertz, S., et al., Geographical location influences the composition of the gut microbiota in wild house mice (Mus musculus domesticus) at a fine spatial scale. PloS one, 2019. 14(9): p. e0222501.

62. Weldon, L., et al., The gut microbiota of wild mice. PLoS One, 2015. 10(8): p. e0134643.

63. Hufeldt, M.R., et al., Variation in the gut microbiota of laboratory mice is related to both genetic and environmental factors. Comparative medicine, 2010. 60(5): p. 336–347.

64. Runge, S., et al., Laboratory mice engrafted with natural gut microbiota possess a wildling-like phenotype. Nature Communications, 2025. 16(1): p. 1–14.

65. Yeung, F., et al., Altered immunity of laboratory mice in the natural environment is associated with fungal colonization. Cell Host & Microbe, 2020. 27(5): p. 809–822. e6.

66. Jameson, S.C. and D. Masopust, Understanding subset diversity in T cell memory. Immunity, 2018. 48(2): p. 214–226.

67. Stegelmeier, A.A., et al., Myeloid cells during viral infections and inflammation. Viruses, 2019. 11(2): p. 168.

68. Tan, W., et al., IL-17F/IL-17R interaction stimulates granulopoiesis in mice. Experimental hematology, 2008. 36(11): p. 1417–1427.

69. Paudel, S., et al., Regulation of emergency granulopoiesis during infection. Frontiers in immunology, 2022. 13: p. 961601.

70. Chen, Q., et al., Sperm tsRNAs contribute to intergenerational inheritance of an acquired metabolic disorder. Science, 2016. 351(6271): p. 397–400.

71. Rosshart, S.P., et al., Wild mouse gut microbiota promotes host fitness and improves disease resistance. Cell, 2017. 171(5): p. 1015–1028. e13.

